# YhcB coordinates peptidoglycan and LPS biogenesis with phospholipid synthesis during *Escherichia coli* cell growth

**DOI:** 10.1101/2021.04.16.440158

**Authors:** Emily C. A. Goodall, Georgia L. Isom, Jessica L. Rooke, Christopher Icke, Karthik Pullela, Bing Zhang, James Rae, Wee Boon Tan, Matthias Winkle, Antoine Delhaye, Eva Heinz, Jean-Francois Collet, Adam F. Cunningham, Mark A. Blaskovich, Robert G. Parton, Jeff A. Cole, Shu-Sin Chng, Waldemar Vollmer, Jack A. Bryant, Ian R. Henderson

**Affiliations:** Institute for Molecular Bioscience, The University of Queensland, St. Lucia, Australia; Institute of Microbiology and Infection, University of Birmingham, UK; Department of Chemistry, National University of Singapore, Singapore; Centre for Bacterial Cell Biology, Biosciences Institute, Newcastle University, Newcastle upon Tyne, UK; de Duve Institute, Université Catholique de Louvain, Brussels, Belgium; Departments of Vector Biology and Clinical Sciences, Liverpool School of Tropical Medicine, Liverpool, UK; Centre for Microscopy and Microanalysis, The University of Queensland, St. Lucia, Australia

**Keywords:** YhcB, cell envelope, polymyxin B, *Escherichia coli*, LPS, peptidoglycan, phospholipid membrane, TraDIS, TIS, transposon mutagenesis

## Abstract

The cell envelope is essential for viability in all kingdoms of life. It retains enzymes and substrates within a confined space while providing a protective barrier to the external environment. Destabilising the envelope of bacterial pathogens is a common strategy employed by antimicrobial treatment. However, even in one of the most well studied organisms, *Escherichia coli*, there remain gaps in our understanding of how the synthesis of the successive layers of the cell envelope are coordinated during growth and cell division. Here, we used a whole genome phenotypic screen to identify mutants with a defective cell envelope. We report that loss of *yhcB*, a conserved gene of unknown function, results in loss of envelope stability, increased cell permeability and dysregulated control of cell size. Using whole genome transposon mutagenesis strategies we report the complete genetic interaction network of *yhcB*, revealing all genes with a synthetic negative and a synthetic positive relationship. These genes include those previously reported to have a role in cell envelope biogenesis. Surprisingly, we identified genes previously annotated as essential that became non-essential in a Δ*yhcB* background. Subsequent analyses suggest that YhcB sits at the junction of several envelope biosynthetic pathways coordinating the spatiotemporal growth of the cell, highlighting YhcB as an as yet unexplored antimicrobial target.

## Introduction

The bacterial cell envelope plays a fundamental role in protection, host interaction, energy generation, expulsion of toxic substances and coordination of growth and cell division. As a physical barrier, it has a central role in acquired and intrinsic antimicrobial resistance. The cell envelope biogenesis systems are therefore important drug targets. In Gram-negative bacteria, the tripartite cell envelope is composed of the cytoplasmic membrane (CM), peptidoglycan (PG) layer and an outer membrane (OM) (Silhavy et al. 2010). The OM is an asymmetrical bilayer of phospholipids and lipopolysaccharide (LPS), which forms a strong permeability barrier conferring resistance to many toxic antimicrobials. Both membranes are studded with integral membrane proteins and peripheral lipoproteins that facilitate cellular functions (Lugtenberg 1981; Molloy et al. 2000; Luirink et al. 2012). Each component of the cell envelope must be synthesised and assembled in a coordinated fashion to maintain cell envelope homeostasis and viability. Thus, understanding how this complex envelope is synthesised and maintained is instrumental to understanding how to disrupt its function, to kill the bacterium, or render the organism susceptible to otherwise ineffective treatments.

Over the last 50 years a variety of sophisticated and complex multiprotein machineries have been discovered that are required for the synthesis of the Gramnegative cell envelope. The Lpt machinery facilitates insertion of LPS into the OM (Sperandeo et al. 2008; Polissi and Sperandeo 2014; May et al. 2015; Simpson et al. 2015; Okuda et al. 2016; Sherman et al. 2018), the BAM complex coordinates OM protein assembly and insertion (Ruiz et al. 2005; Wu et al. 2005; Knowles et al. 2009), the Lol system traffics lipoproteins across the periplasm for incorporation into the OM (Matsuyama et al. 1995; Matsuyama et al. 1997; Yakushi et al. 2000; Okuda and Tokuda 2011), the Mla complex enables phospholipids transport between the two membranes (Malinverni and Silhavy 2009; Thong et al. 2016; Abellón-Ruiz et al. 2017; Powers and Trent 2018; Ercan et al. 2019; Hughes et al. 2019; Shrivastava and Chng 2019; Chi et al. 2020), and the elongasome and divisome are the architects of peptidoglycan assembly during cell elongation and division (Goehring and Beckwith 2005; Den Blaauwen et al. 2008). Each pathway in isolation is broadly understood. However, the precise molecular events that govern the crosstalk between the different biosynthetic pathways, both spatially and temporally during growth and cell division, remain to be elucidated.

Genetic screens to identify synthetically lethal interactions and suppressor mutations have played a significant role in identifying the interconnected pathways of envelope biosynthesis (Ogura et al. 1999; Klein et al. 2014). Transposon insertion sequencing (TIS) is one approach that enables the study of genetic interactions on a whole genome scale (DeJesus *et al*. 2017). Here we used a chemical genomics approach to identify genes required for cell envelope homeostasis in *Escherichia coli*, coupling TraDIS, a TIS method (Langridge et al. 2009; Cain et al. 2020), with the membrane targeting antibiotic polymyxin B. We demonstrate that a poorly characterised protein, YhcB, is crucial for tolerance to polymyxin B and for maintenance of cell envelope homeostasis in *E. coli*. Previous studies revealed that YhcB is a CM protein widely conserved in Gammaproteobacteria that physically interacts with the elongasome, the machine that coordinates peptidoglycan synthesis along the cylindrical part of the cell, and with LapA, a protein involved in regulating LPS biosynthesis (Mogi et al. 2006; Den Blaauwen et al. 2008; Maddalo et al. 2011; Li et al. 2012; Klein et al. 2014; Mahalakshmi et al. 2014; Fivenson and Bernhardt 2020). We demonstrate that loss of YhcB results in dysregulation of cell length and width. We created a high-density transposon mutant library in an *E. coli* Δ*yhcB* strain to gain a whole genome view of genetic interactions with *yhcB*. Subsequent screening of this library against chemical stresses revealed all genes that suppress the envelope defects associated with the loss of *yhcB*. These data suggest that YhcB is stationed at the interface between PG and LPS synthesis, and phospholipid membrane biogenesis, and has a role in coordinating the spatial and temporal assembly of the cell envelope.

## Results

### Identification of envelope barrier-defective mutants

Previously, we described a high-density transposon-mutant library of *E. coli* K-12 (Goodall et al. 2018). To identify genes required for cell envelope biogenesis we applied a chemical genomics approach exposing our library to sub-inhibitory concentrations of the membrane-acting antibiotic polymyxin B (Fig. S1A). Polymyxins bind the lipid A moiety of LPS, displacing divalent cations, disrupting the integrity of LPS-crosslinks, and resulting in an increase in membrane permeability (Newton 1953; Dixon and Chopra 1986; Trimble et al. 2016; Li and Velkov 2019). We posited that if the transposon disrupts a gene required for the maintenance of cell envelope integrity, the resulting mutant will become more susceptible to polymyxin B, and the gene would be identified through our TIS screen as these mutants will be outcompeted during growth; identifiable as a depletion in transposon-insertions. We grew the transposon library in LB broth with or without 0.2 μg/ml polymyxin B, in duplicate, and harvested cells after ~5 generations of growth (Fig. 1A). The sequencing yielded a total of >4.2 M reads, estimated to sample >99% of the possible unique insertion sites (Fig. S1B). We used the BioTraDIS pipeline for data processing and analysis (Barquist et al. 2016), and mapped the processed reads to the *E. coli* BW25113 reference genome obtained from NCBI (accession CP009273.1). The insertion index scores (IISs; the number of insertions per gene, normalised for gene length) between replicates were highly comparable (Fig. S1C). We identified 54 genes required for growth in sub-inhibitory concentrations of polymyxin B (Fig. 1B; Table S1). Comparison of the relative abundance of COG (Cluster of Orthologous Groups) categories of these 54 genes showed a marked increase in genes involved in cell wall/membrane/envelope biogenesis (COG category ‘M’), supporting the validity of the TIS screen (Fig. S1D). Many of these genes (e.g. *bamB, degP, galE, surA*, and *waaBCFGJPR*) have been identified in a similar TIS screen using *Klebsiella pneumoniae* or have previously been reported as mutants sensitive to polymyxin (Yethon et al. 2000; Jana et al. 2017). The COG analysis identified only three genes (*ydbH, yqjA* and *yhcB*) of unknown function (category ‘S’ in Fig. S1D) that were important for survival when exposed to sub-inhibitory concentrations of polymyxin B (Fig. 1C and Fig. S1E). YdbH is a putative autotransporter, while YqjA belongs to the DedA family of proteins that have been suggested to be involved with membrane homeostasis (Keseler et al. 2017). Given that *yhcB* is conserved in three out of the six ESKAPE pathogens, is required for tolerance to polymyxin, an antibiotic of last resort, and is the target of a sRNA toxin that causes cell death, we decided to investigate the role of YhcB further (Mogi et al. 2006; Choi et al. 2018).

**Figure 1.**
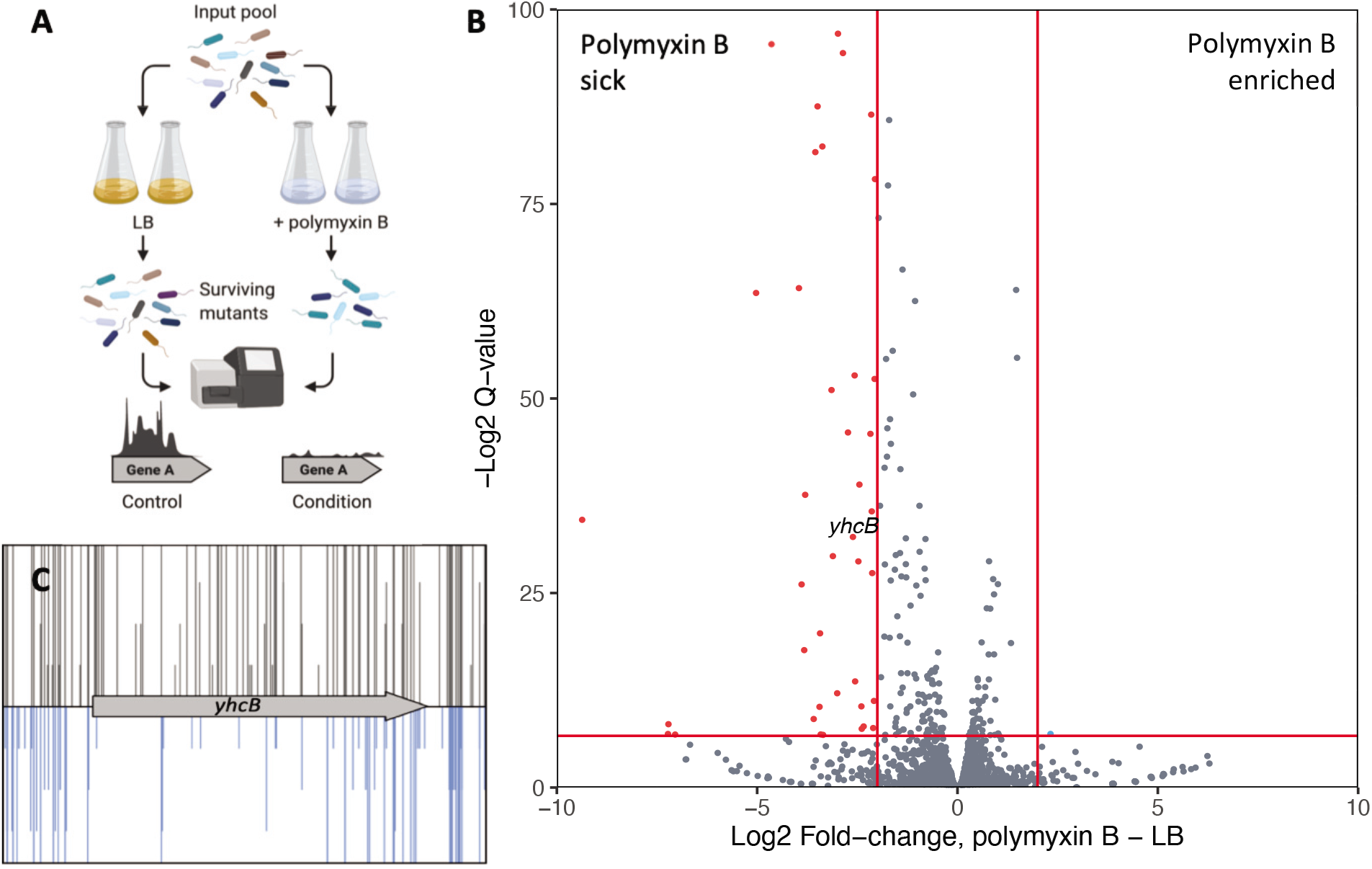
Screening the BW25113 transposon library in sub-inhibitory concentrations of polymyxin B. (A) Schematic showing the TIS experiment for identification of polymyxin B-sensitive mutants. (B) Volcano plot of the fold-change in mapped reads between conditions, calculated using BioTraDIS. Datapoints in red correspond with genes with >2-fold decrease in reads in the polymyxin B dataset relative to the LB outgrowth dataset, and with a Q-value > 0.01 (red horizontal line). While one datapoint in blue had a >2-fold increase in reads in polymyxin B dataset relative to the control, and a Q-value > 0.01. A label for the *yhcB* datapoint has been added above the point for clarity. (C) Image of the *yhcB* insertion data following outgrowth in LB only (grey, above) or in LB supplemented with polymyxin B (blue, below). The transposon insertion position along the gene is marked by a vertical line, with the vertical line size corresponding with read depth, with visibly fewer transposon-insertion sites identified within *yhcB* following outgrowth in LB supplemented with polymyxin B.

### *Deletion of* yhcB *confirms an envelope defect*

To confirm the barrier defect of the *yhcB* mutant, we grew the parent strain, *E. coli* BW25113, and an isogenic Δ*yhcB* mutant on LB agar plates supplemented with either vancomycin, a high molecular weight antibiotic that is typically unable to cross the Gramnegative OM, or plates supplemented with SDS and EDTA (Nikaido and Vaara 1985; Nikaido 2005). The Δ*yhcB* mutant was more sensitive than the parent strain to both conditions (Fig. 2A). Ectopic expression of YhcB under arabinose induction on a pBAD plasmid was able to complement sensitivity defects, consistent with previous reports (Sung et al. 2020). These data confirm that loss of *yhcB* results in a severe envelope defect (Fig. S2A).

**Figure 2.**
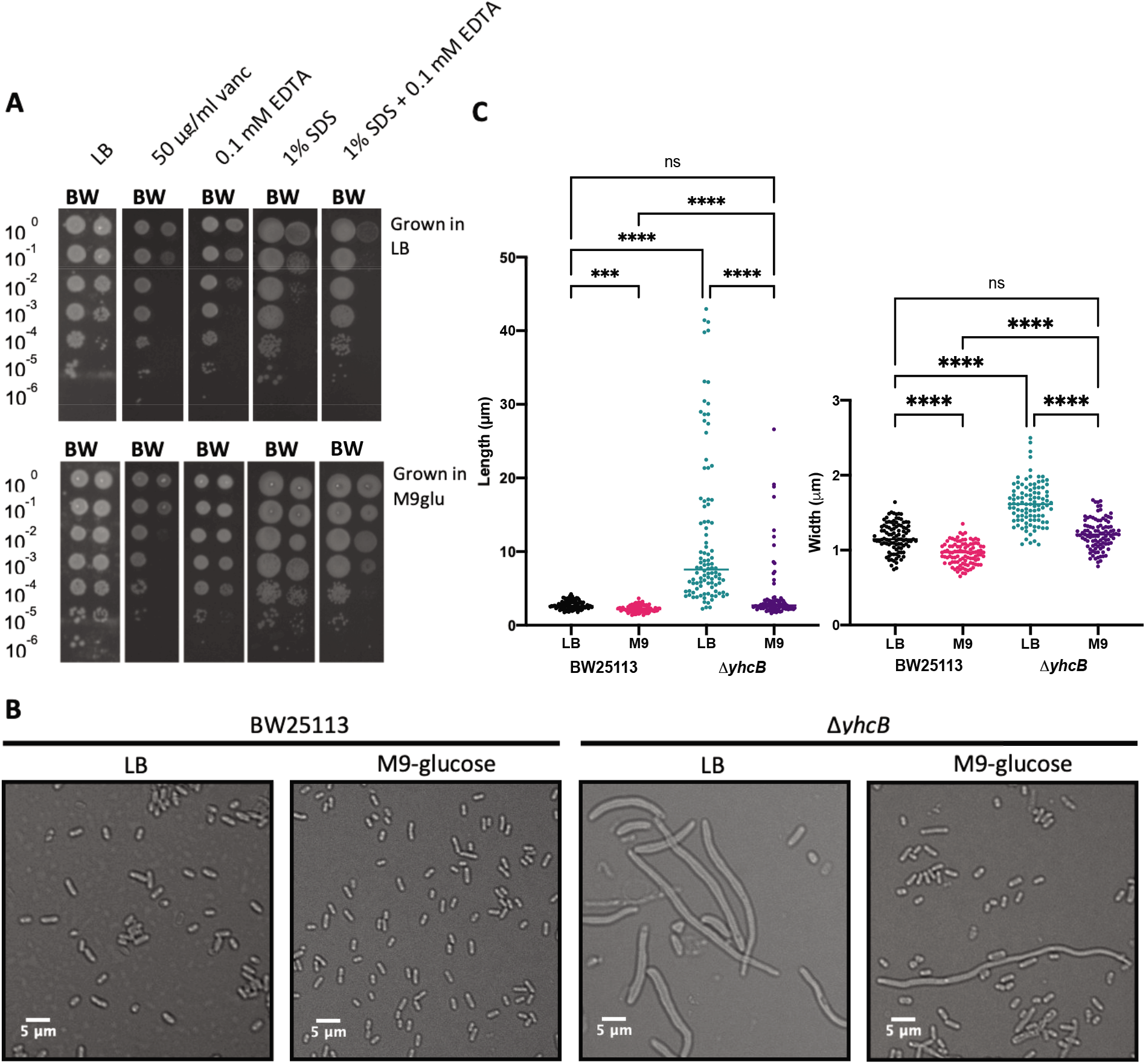
Validation of a *yhcB* mutant cell envelope defect and the effect of growth medium on cell size. (A) Overnight cultures of the WT strain BW25113 (BW) and Δ*yhcB* (Y) strain grown in LB or M9-glucose at 37°C, normalised to an OD_600_ of 1.00 and 10-fold serially diluted before inoculating LB agar plates supplemented with various compounds. (B) The WT and Δ*yhcB* mutant were grown overnight in LB or M9-glucose media at 37°C. Cells were visualised using a Nikon 90i microscope, a 5 μm bar is shown for scale. (C) Cell measurements for BW25113 and Δ*yhcB* mutant grown in LB and M9-glucose. Images were taken after overnight growth and are representative of n=4 experiments, data from 1 experiment is presented. Width and Length measurements were taken of 100 cells in each condition. A Kruskal-Wallis test with Dunn’s correction for multiple comparisons was used to compare between samples: p < 0.0001 (****); p < 0.0002 (***).

### Effect of YhcB on cell size

Despite normalising cultures by optical density, the number of viable colony forming units (CFUs) of the Δ*yhcB* mutant were 10-fold fewer than the parent on the LB control plate (Fig. 2A). Analysis of cellular morphology by DIC microscopy revealed that the Δ*yhcB* mutant cells were significantly larger than the parent strain when grown in LB (Fig. 2B). Expression of *yhcB* in the Δ*yhcB* mutant background substantially decreased the cell dimensions relative to the Δ*yhcB*-pBAD empty vector strain but did not restore cell size completely (Fig S2B). We also observed that the ectopic expression of *yhcB* in the *E. coli* BW25113 parent strain resulted in longer cells than the control. Taken together, these data suggest that the abundance of YhcB within the cell impacts cellular dimensions, and that YhcB is important for maintaining cell size and shape.

Bacterial cell dimensions are influenced by a number of factors (reviewed by Cesar and Huang 2017), but cell size is fundamentally determined by growth rate and nutrient availability (Schaechter et al. 1958; Hill et al. 2013; Vadia and Levin 2015; Vadia et al. 2017). A shift from growth in nutrient-restricted to nutrient-rich medium results in an increase in both cell length and cell width (Pierucci 1978). Therefore, we investigated the effect of growth medium on the size of the Δ*yhcB* mutant. Overnight growth in M9 minimal medium supplemented with glucose (M9-glucose) substantially decreased the cell size of the Δ*yhcB* mutant compared to growth in LB, although a small subset of cells within the population still exhibited a division defect and increased cell length (Fig. 2B-C). Moreover, overnight growth in M9-glucose restored resistance to membrane-acting compounds SDS and EDTA, but cells remained susceptible to vancomycin (Fig. 2A). Changes in temperature alter growth rate but not size (Schaechter et al. 1958). To understand whether the fitness advantage conferred by growth in M9-glucose was due to a slower doubling time, or nutrient-dependent, we grew the Δ*yhcB* mutant in a range of conditions. Overnight growth in M9-glucose and M9-glycerol at 37°C, and LB at 16°C, restored resistance to both SDS and EDTA, but no growth condition conferred resistance to vancomycin (Fig. S3). Altogether these results indicate that a slower growth rate partially alleviates the defect caused by deletion of *yhcB* but does not restore the envelope permeability barrier.

We hypothesised that there is an envelope synthesis defect in a Δ*yhcB* mutant that becomes more severe over time during growth in LB. To investigate this, overnight cultures (grown first in M9 medium) were sub-cultured in LB at a starting optical density (OD) of 0.05. Bacteria were grown at 37°C with aeration, and both the OD_600_ and the corresponding number of CFUs were recorded hourly. The rate of increase in biomass of the mutant, as assessed by optical density, was comparable to that of *E. coli* BW25113 (Fig. 3). In contrast, upon transition to stationary phase, the number of Δ*yhcB* CFUs decreased 10-fold from ~3.5 x 10^8^ to ~3.5 x 10^7^. These data suggest that the fast-growing mutant cells gradually expand in cell size, fail to divide, but lose viability when they enter the stationary growth phase.

**Figure 3.**
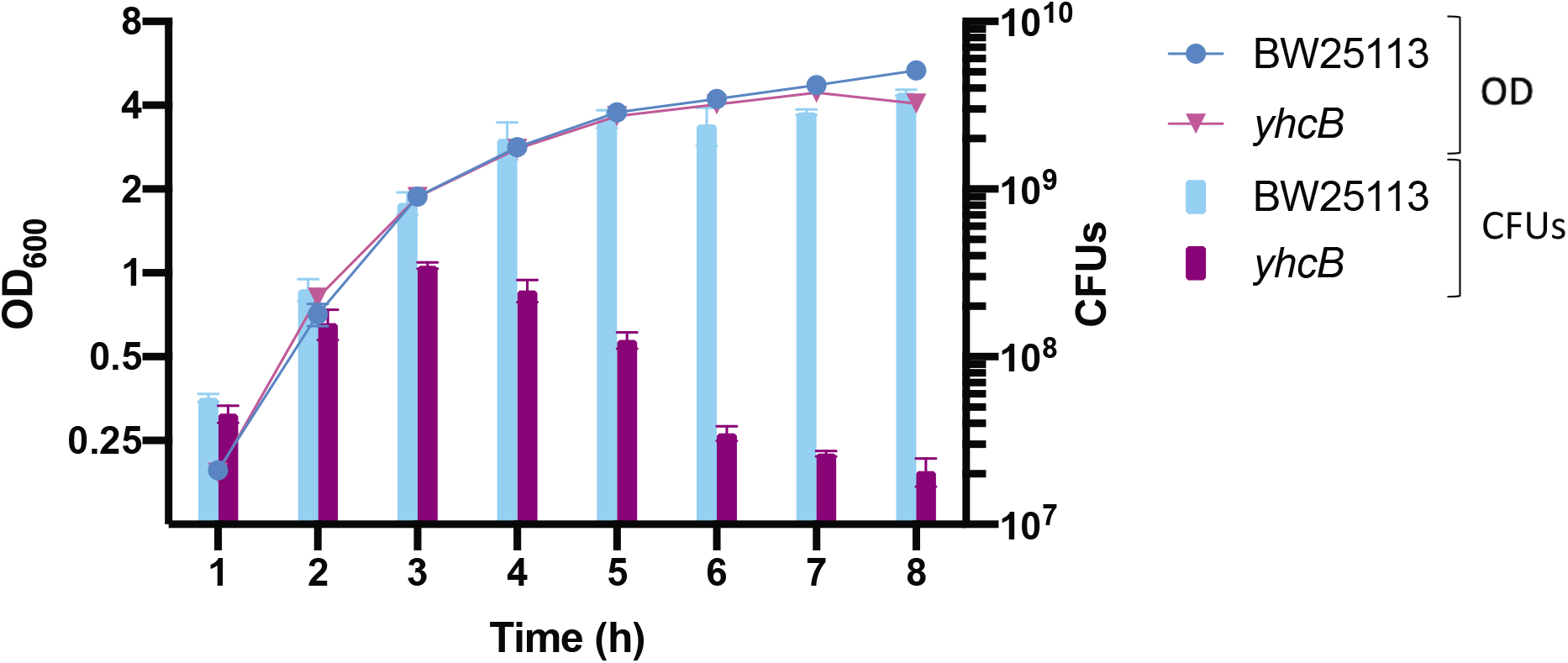
The effect of *yhcB*-deletion during growth in rich media. (A) The optical density (OD) and number of colony forming units (CFUs) were recorded hourly over 8 h during growth in LB at 37°C. Strains were grown in triplicate, the mean is plotted and the s.d. represented by error bars. BW25113 is shown in blue (•), *yhcB* is shown in pink (▾).

### The domains of YhcB contribute to survival under different conditions

To gain a better understanding of how YhcB functions, we scrutinised the primary structure of the protein. A prediction of YhcB secondary structure of YhcB by PSIPRED supports the model put forward by Mogi et al. (2006; Jones 1999; Buchan and Jones 2019). YhcB has a single transmembrane (TM) domain, with a cytoplasmic helical domain and a disordered cytoplasmic C-terminal region (Fig. S4A); the topology of which has been confirmed by GFP- and PhoA-fusion analysis (Maddalo et al. 2011). A multisequence alignment of the amino acid sequence of YhcB from 150 different species was generated using the online program ConSurf (Ashkenazy et al. 2016). This revealed two highly conserved amino acid motifs within the disordered C-terminus: a ‘NPF’ motif after the long cytoplasmic helical domain, followed by a non-conserved linker region, and a highly conserved ‘PRDY’ motif at the extreme C-terminus (Fig S4B). A large number of transposon insertions were identified within the 3’ end of *yhcB* in the initial polymyxin TIS screen; the *yhcB* gene only met the stringency criteria for identification when 20% of the 3’ end of the coding sequence (CDS) was discarded. We mapped the coordinates of these domains to the original TIS data and the data suggest that the ‘PRDY’ domain is dispensable for survival in the presence of polymyxin B (Fig. 4A). To confirm the functional importance of the ‘PRDY’ domain, which is highly conserved (Fig. S5), we constructed complementation vectors encoding truncated variants of YhcB and repeated our plate-based envelope screens (Fig. 4B). Consistent with results reported by Sung *et al*. (2020), the ATM construct fully complemented both phenotypes. However, deletion of the ‘PRDY’ domain rendered the mutant sensitive to vancomycin, but not SDS-EDTA (Fig. 4C). These data suggest a dual function of the YhcB protein: the cytoplasmic helical domain is needed to suppress sensitivity to SDS and EDTA, while the function of the ‘PRDY’ domain is needed for survival in the presence of vancomycin, but not polymyxin B or SDS and EDTA.

**Figure 4.**
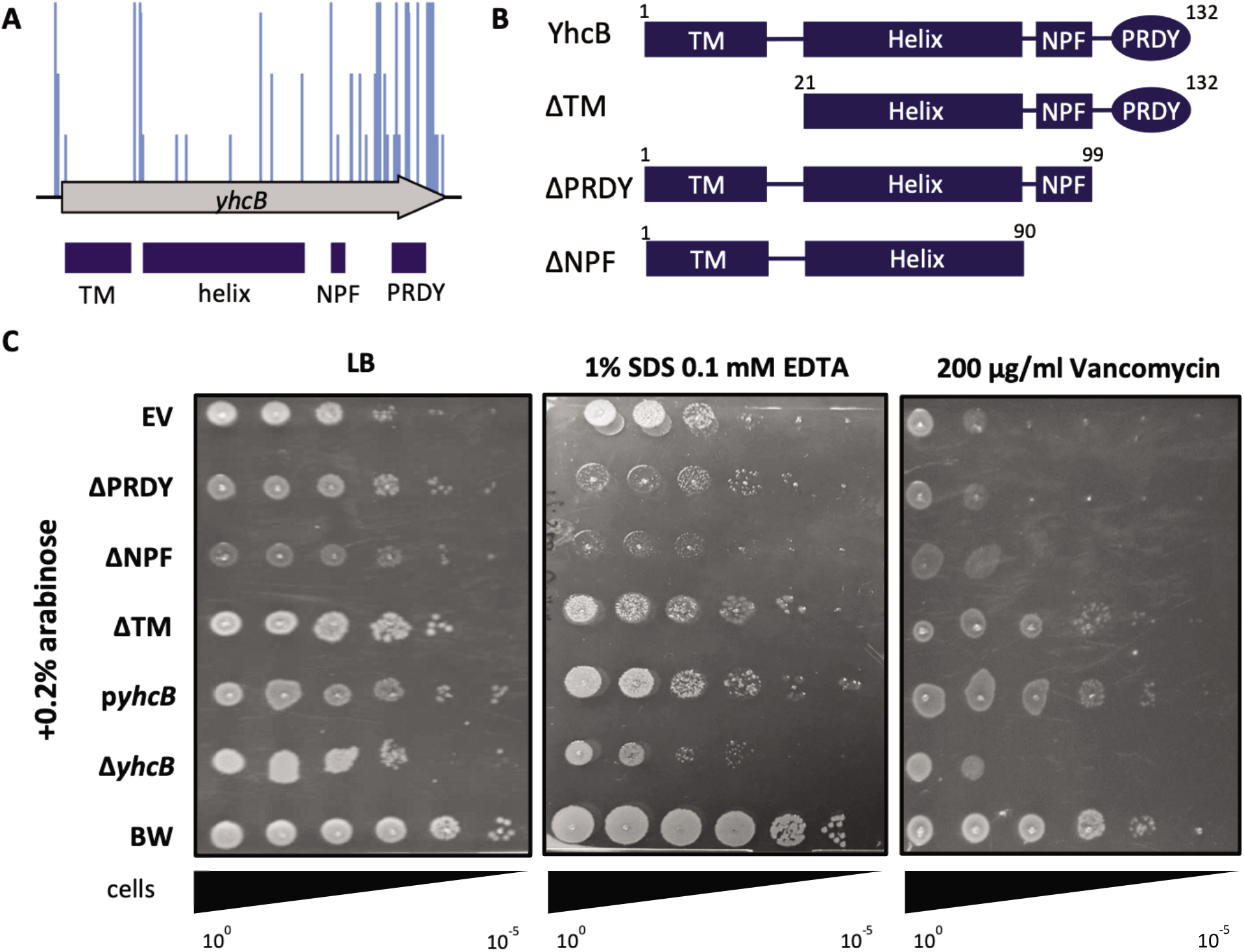
The structural and functional contribution of the domains of YhcB. (A) Polymyxin B TIS screen data (blue insertions) with the domains of YhcB mapped underneath (dark blue boxes). (B) Schematics of the plasmid-based YhcB complementation constructs. (C) Overnight cultures grown in LB supplemented with carbenicillin (for plasmid selection), normalised to an OD_600_ of 1.00 and 10-fold serially diluted before inoculating LB agar plates supplemented with vancomycin or SDS and EDTA, with 0.2% arabinose.

### *Identification of genetic interactions with* yhcB

To identify all genes with a synthetically lethal relationship with *yhcB*, a transposon mutant library was constructed in a Δ*yhcB* background using a mini Tn5 transposon encoding a kanamycin resistance gene. A second library was constructed in the *E. coli* BW25113 parent strain using the same transposon as a control. Approximately 800,000 transposon-mutants were collected and pooled for each library. Two technical replicates of each library were sequenced (Fig. S6A; Table S2). There was a high correlation coefficient between the gene insertion index scores of each replicate for both libraries (R^2^ = 0.93 and R^2^ = 0.96 for the WT and Δ*yhcB* library respectively). The insertion sites were evenly distributed around the genome (Fig. 5A and Fig. S6B), and the respective insertion density amounted to an average of one insertion every 6.28 bp in the WT library, and one insertion every 6.99 bp in the Δ*yhcB* library. We quantified insertions per CDS to identify genes that were sparsely disrupted by transposon insertion events i.e. essential genes in each genetic background (Barquist et al. 2016). There was an overlap of 382 essential genes between the two datasets and these were not considered further (Fig. 5B).

**Figure 5.**
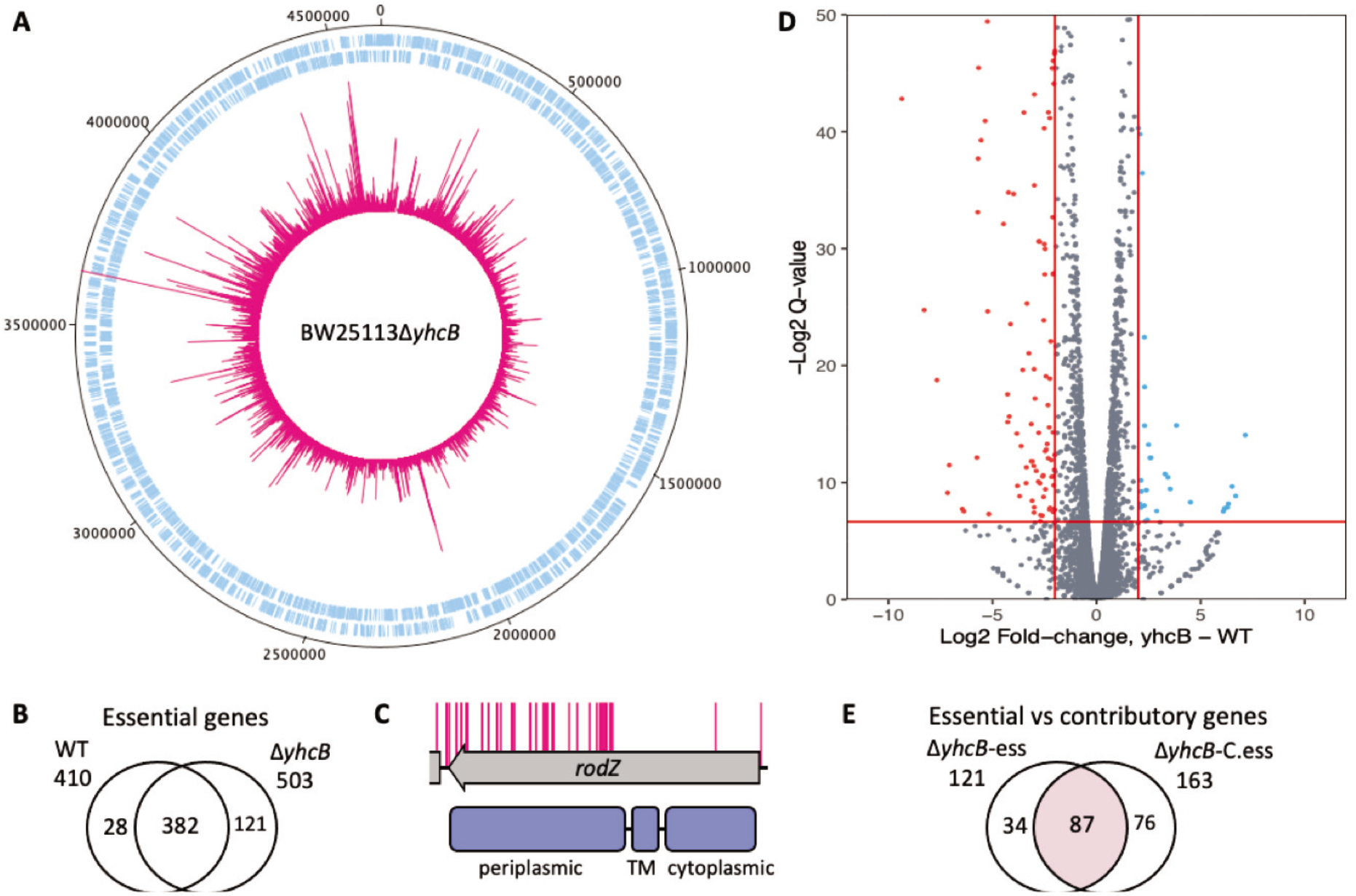
Construction of a transposon library in a *yhcB* mutant and identification of genes with a synthetic lethal relationship with *yhcB*. (A) The Δ*yhcB* transposon library. A genome map of BW25113 starting at the annotation origin with the sense and antisense coding sequences of BW25113 shown in blue, respectively, and the position and frequency of sequenced transposon insertion events shown in pink. (B) Comparison of genes identified by bimodal analysis as significantly underrepresented by transposon mutants in each library. (C) The insertion profile of *rodZ* in the Δ*yhcB* library, with RodZ domains shown below in blue. (D) The log_2_ fold-change of read depth per gene between the parent BW25113 and the Δ*yhcB* transposon library. Genes enriched for transposon insertions (>2-fold) in the *yhcB* library are shown in blue, indicative of more fit mutants. Genes with a >2-fold decrease in insertions are shown in red, indicative of sick mutants. A Q-value threshold of 0.01 was used (red horizontal line) (E) Comparison of genes predicted to be essential (sparsely disrupted genes) with genes identified as having a >2-fold decrease in reads compared to the control library (genes that contribute to fitness). This is a comparison of the bi-modal and enrichment analyses outputs. The overlap of these analyses are the 87 identified synthetic lethal genes (pink).

Previously, Li *et al*. (2012) reported that the deletion of both *rodZ* and *yhcB* in combination is lethal, genetically linking the elongasome with the function of YhcB. Our data confirmed a synthetic lethal interaction between *rodZ* and *yhcB*. However, we observed that only the 5’ end of the *rodZ* gene, corresponding with the cytoplasmic helixturn-helix and transmembrane domains of the RodZ protein, is essential in a *yhcB* background (Fig. 5C). These domains are required for maintaining cell shape (Shiomi et al. 2008; Bendezú et al. 2009). This result supports the validity of our approach. Surprisingly, our bimodal analysis of the data revealed 28 genes that were predicted to be essential in a WT library but no longer essential when *yhcB* is deleted (Fig. 5B, Table S3). This list includes *ftsH* (Fig. S6C), which encodes a metalloprotease that degrades several proteins, including LpxC. Deletion of FtsH results in toxic accumulation of LPS via increased LpxC stability, which is lethal to the cell (Ogura et al. 1999). The increased number of transposons in *ftsH* in the Δ*yhcB* library suggest that an increase in LpxC stability in this background is less toxic.

The bimodal analysis revealed 121 genes that were predicted to be essential in a Δ*yhcB* background. However, this analysis pipeline does not identify the non-essential genes that contribute to fitness: such mutants are viable but either more or less fit. Consequently, when compared to the control library they are represented by higher or lower numbers of sequencing reads, respectively. Therefore we further analysed our data using the tradis_comparison.R script (Barquist et al. 2016). This analysis uses edgeR and measures the fold-change of sequence read depth between a condition and control (Robinson et al. 2009). The 382 genes reported to be essential in both libraries were removed from this analysis. After filtering our data, this method identified 22 genes that, when disrupted, confer a fitness advantage (Fig.5D, blue), and 163 genes that, when disrupted, confer a fitness defect in a Δ*yhcB* mutant (Fig. 5D, red; Table S4; Fold change > 2, Q-value < 0.01). By combining these analyses, we defined synthetically lethal genes as those that are both significantly underrepresented by transposon insertion mutants and that have a >2-fold decrease in reads when compared to the control with a Q-value < 0.01. Altogether 87 genes met these criteria: we consider these genes to be synthetically lethal with *yhcB* and they are discussed below (Fig. 5E).

### *Identification of pathways important for viability of a* yhcB *mutant*

As YhcB interacts directly with components of the elongasome, and is synthetically lethal with *rodZ*, we inspected the list of synthetic lethal genes for additional genes involved in peptidoglycan synthesis, remodelling or recycling. Five genes (*mepM, mepS, dacA, dapF* and *ldcA*) were identified (Fig. 6A). The genes *mepM* and *mepS* encode two of the three peptidoglycan endopeptidases that cleave the 3-4 *meso*-Dap-d-Ala crosslink and are collectively essential for cell elongation in *E. coli* (Singh et al. 2012). However, a third endopeptidase of this group encoded by *mepH*, was not essential in a Δ*yhcB* mutant (Fig. 6A-B). DapF is an epimerase that catalyses the conversion of l,l-diaminopimelate (LL-DAP) to *meso*-diaminopimelate (*meso*-DAP), which is an integral component of the peptidoglycan stem peptide and the primary residue from which cross-links are formed (Richaud et al. 1987). Sacculi isolated from *dapF* mutants have fewer crosslinks (Mengin-Lecreulx et al. 1988). The DD-carboxypeptidase PBP5, encoded by *dacA*, removes the terminal D-alanine from peptidoglycan pentapeptides (Spratt and Strominger 1976), and is the primary carboxypeptidase under standard laboratory growth conditions (Nelson and Young 2000; Nelson and Young 2001). Finally, the cytosolic LD-carboxypeptidase LdcA participates in the recycling of peptidoglycan turnover products (Fig. 6A). Deletion of *ldcA* results in the lysis of cells during entry into the stationary growth phase due to the accumulation of peptidoglycan crosslinking defects (Templin et al. 1999). From these data, it could be inferred that a Δ*yhcB* mutant cannot survive when additional mutations weaken the integrity of the peptidoglycan layer.

**Figure 6.**
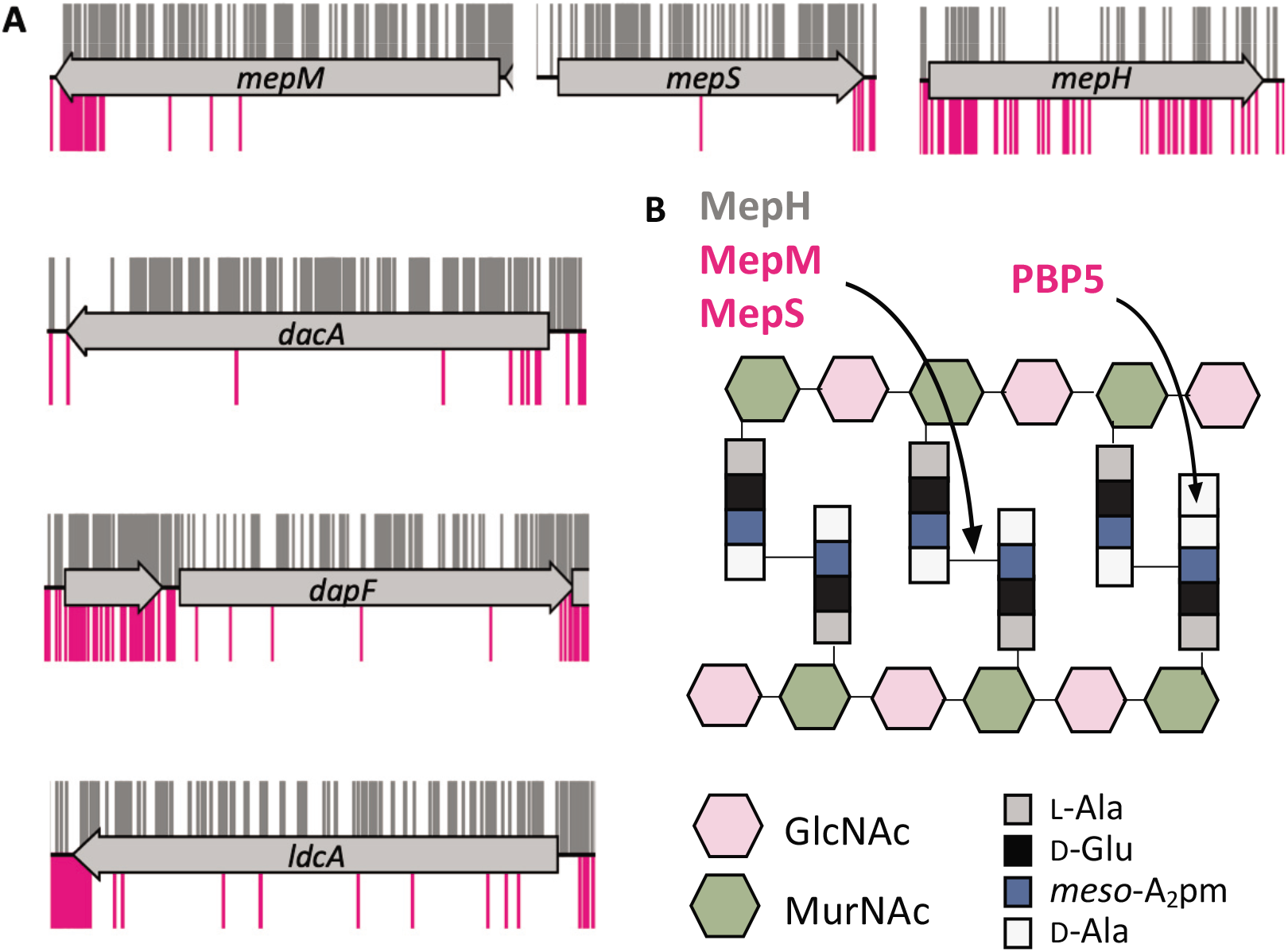
*yhcB* is synthetically lethal with components of PG synthesis and recycling pathways. (A) TIS data of the BW25113 library (grey) and BW25113Δ*yhcB* library (pink) were plotted above and below the gene track, respectively. Insertion sites are represented by vertical bars and are capped at a frequency of 1. (B) Schematic representation of the target sites of the endopeptidases and PBP5 (encoded by *dacA*). Abbreviations: *meso*-diaminopimelate (*meso*-A_2_pm); N-acetylglucosamine (GIcNAc); N-acetylmuramic acid (MurNAc); L-Ala, L-alanine; D-Glu, D-glutamic acid; D-Ala, D-alanine.

To determine whether other cellular processes or pathways are enriched among the genes identified as synthetically lethal with *yhcB*, we used the PANTHER overrepresentation test (Mi et al. 2021). This analysis compares the identified proportion of genes in a given functional category to the expected number of genes in a pathway derived from whole genome data (Table S6). Four processes were identified as functionally enriched among the list of synthetic lethal genes: the ‘enterobacterial common antigen (ECA) biosynthetic pathway’, ‘LPS core biosynthesis’, ‘cell division’, and ‘response to abiotic stimulus’ (Fig. 7A). With the exception of the *tol-pal* system (Fig. S7A), the remaining genes listed within the ‘cell division’ and ‘response to abiotic stimulus’ categories did not share overlapping functions and we did not pursue these further.

**Figure 7.**
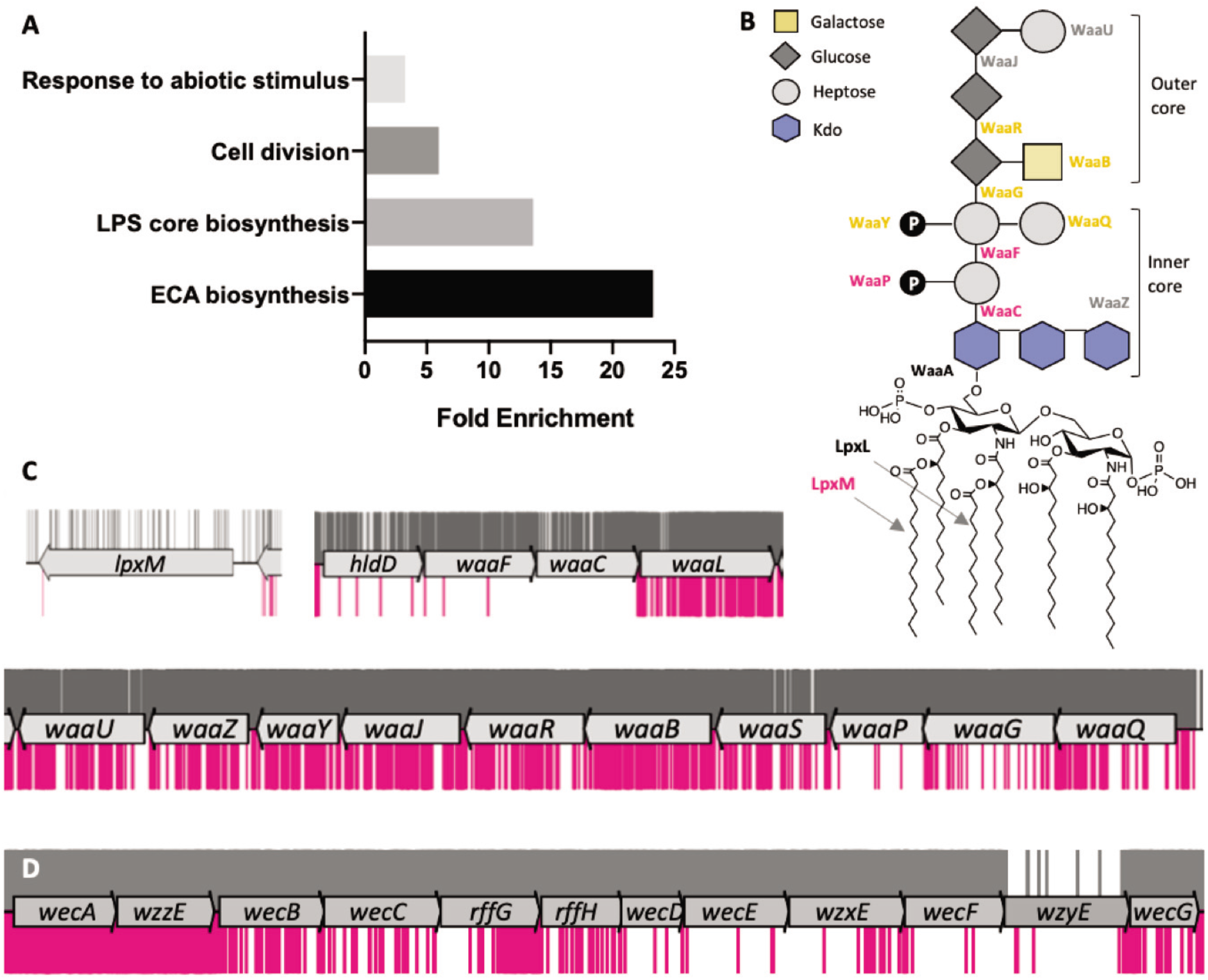
Functional enrichment among synthetic lethal genes. (A) Functional enrichment of cellular processes among the genes that are synthetically lethal with *yhcB*. Fisher’s exact test with Bonferroni correction for multiple testing. Results displayed for P < 0.05. (B) A schematic of the structure of LPS. LPS biosynthesis enzymes are indicated next to the linkage they form (central column) or component they ligate (side branches). Synthetic lethal enzymes are labelled in dark pink, enzymes that contribute to fitness are labelled in orange, core essential enzymes in black and nonessential enzymes in grey. (C) TIS data for BW25113 (grey) and BW25113Δ*yhcB* (pink) libraries were plotted above and below the gene track, respectively. Insertion sites are represented by vertical bars and are capped at a frequency of 1. Insertions within genes required for LPS core biosynthesis, and *lpxM*, are significantly underrepresented. (D) The insertion profiles in genes of the enterobacterial common antigen biosynthetic pathway. Abbreviations: Lipopolysaccharide (LPS); Enterobacterial Common Antigen (ECA).

Upon closer inspection of LPS biosynthetic pathways, both the heptosyl transferases WaaC and WaaF, and the entire heptose biosynthetic pathway were synthetically lethal (Fig. 7B-C, Fig. S7B). The WaaP kinase, which phosphorylates the first heptose of the inner core, was also essential in a *yhcB* mutant. Enzymes WaaBGQRY were not essential in a Δ*yhcB* background, but mutants were less fit. In contrast, ligation of the third Kdo moiety by WaaZ was non-essential. The acylation of lipid A by LpxM was also essential in a Δ*yhcB* strain (Fig. 7C). Together these data suggest that LPS stability, whether mediated by Lipid-A hexa-acylation or LPS crosslinking interactions, is important for viability of a Δ*yhcB* strain.

Within the locus required for ECA biosynthesis the genes *wecCDEF, rffH* and *wzxE* were synthetically lethal with *yhcB*, while disruption to *wecB* and wecG suggests these genes were not synthetically lethal but the mutants were sick (Fig. 7D, Fig S7C). There were no significant differences in the insertion frequency within genes *wzzE* and *wecA*. WzzE mediates the length of the ECA chains by determining the number of repeating units (Barr et al. 1999), and WecA catalyses the first committed step in ECA biosynthesis: the transfer of *N*-acetylglucosamine-1-phosphate onto undecaprenyl phosphate (Und-P) to form ECA-lipid I. In short, an ECA-null mutant is viable and mutants with variable ECA lengths are also viable; only mid-pathway blocks are lethal. Mutations that introduce a mid-pathway block in ECA biogenesis are known to cause an aberrant cell morphology (Jorgenson *et al*. 2016), due to accumulation of undecaprenyl-linked ECA-lipid II intermediates creating limited availability of Und-P: a compound of limited abundance that sits at the start of several cell envelope biosynthetic pathways (Jorgenson *et al*. 2016). These observations suggest that the combined defects of *yhcB*-deletion and Und-P sequestration are lethal, and indicate that the availability of Und-P might be limited in a *yhcB* mutant.

### *Identification of mutations that suppress a* yhcB-*deletion defect*

In addition to synthetic lethal interactions, mutations that restore a phenotypic defect, so-called suppressor mutations, can assist in the identification of interconnected cellular pathways (Ogura et al. 1999). We applied TIS to identify, at a whole genome scale, all mutations that can restore tolerance to vancomycin or SDS and EDTA, and therefore suppress a Δ*yhcB* phenotype. First, the Δ*yhcB* mutant library was screened on LB agar supplemented with either SDS and EDTA or vancomycin at concentrations that kill the Δ*yhcB* strain. Transposon mutants of the Δ*yhcB* library able to grow under these conditions were identified as before (Fig. 8A-B). We used the recently published AlbaTraDIS package to analyse our data (Page et al. 2020). We identified 28 gene-deletion suppressor mutations shared between both conditions, and therefore considered these to be universal suppressors (Fig. 8C; Table S7). Of note among the 28 universal suppressors was the Mla pathway. All genes of the pathway have insertions along the full length of each CDS indicating a disruption at any stage of the pathway is restorative to both vancomycin and SDS and EDTA sensitivity (Fig. 8D). This was unexpected as *mla* mutants are highly sensitive to SDS and EDTA (Malinverni and Silhavy 2009; Ekiert et al. 2017; Isom et al. 2017).

**Figure 8.**
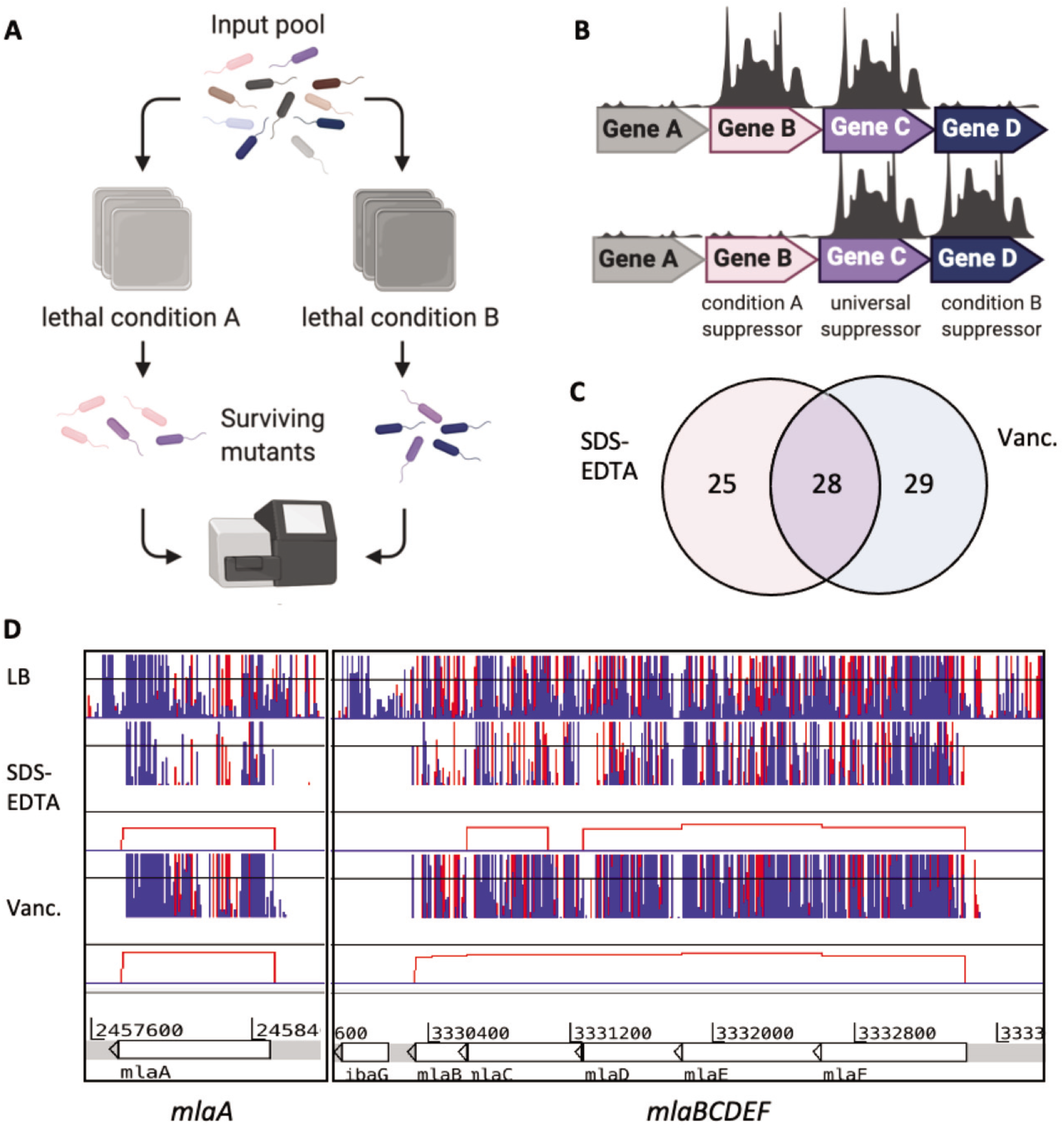
Identification of suppressor mutations that restore a *yhcB*-deletion defect. (A) Experimental overview. (B) Cartoon of output. (C) Overlap between gene-deletion suppressors identified in the lower concentration vancomycin and SDS + EDTA screens. (D) Transposon insertion data for suppressor screens. Red and blue lines indicate the transposon insertion position corresponding with the transposon orientation at the point of insertion. The height of the bar corresponds with mapped sequencing read frequency. The red boxes underneath each suppressor dataset represent significant differential abundance of insertions of the condition compared to the control, identified by AlbaTraDIS. Panel A and B schematics created with BioRender.com, Panel D insertion plots visualised in Artemis (Rutherford et al. 2000).

Deletion of *nlpl* was also identified as a universal suppressor (Fig. S8A). Nlpl is an OM lipoprotein that functions as an adaptor protein for peptidoglycan endopeptidases (Banzhaf *et al*., 2020) and mediates the degradation of MepS by the protease Prc (Tadokoro et al. 2004; Singh et al. 2015). In a Δ*nlpl* strain, MepS activity is significantly increased (Singh *et al*. 2015), which has been shown to enhance peptidoglycan synthesis by stimulating PBP1B-mediated peptidoglycan synthesis and directing peptidoglycan precursors away from the elongasome complex (Lai et al. 2017), presumably to facilitate the repair of defects in the PG (More et al., 2019). Consistent with this hypothesis, *mrcB* (PBP1B) was identified in the list of genes that confer a fitness defect on a *yhcB* mutant strain when deleted (Table S4), while deletion of *mrcA* (PBP1A) and *lpoA*, which encodes the PBP1A activator, were identified as universal suppressors. Suppressor analysis therefore supported a functional link between YhcB and PG synthesis.

Another universal suppressor we identified was deletion of *fabF*, genetically linking *yhcB* with fatty acid biosynthesis (Fig. S8B). FabF, together with FabB, are the two β-ketoacyl-[acyl carrier protein] synthases involved in fatty acid elongation (Garwin et al. 1980a). FabB is the major synthase and is essential while FabF is predominantly required at low temperatures to increase membrane fluidity by increasing the proportion of diunsaturated phospholipids (Garwin et al. 1980b), as FabF is more efficient than FabB at elongating palmitoleic acid (16:1Δ9) to cis-vaccinic acid (18:1Δ11). Therefore, loss of *fabF* might result in decreased membrane fluidity or a decreased rate of phospholipid synthesis, or both.

We plotted the total suppressor insertion data on a genome map to view relative mutant abundance; the assumption here is that read depth is representative of mutant abundance, which correlates with fitness and is an indicator of the strength of the suppression. In addition to Δ*fabF*, disruption of the *mla* operon and Δ*nlpl* we observed a substantial peak at the *uppS-cdsA* locus in both conditions (Fig. 9A). UppS, encoded by *uppS* (*ispU*), is an undecaprenyl pyrophosphate synthase, which synthesises undecaprenyl diphosphate (Und-PP), the only source of *de novo* synthesised Und-PP and the precursor to Und-P (Apfel et al. 1999; Kato et al. 1999). CdsA sits before the branchpoint in the synthesis of the major phospholipids and catalyses the synthesis of cytidine diphosphate-diacylglycerol from phosphatidic acid (Kanfer and Kennedy 1964; Ganong et al. 1980). Both UppS and CdsA are essential, as such, neither gene can be disrupted by transposon insertion. However, we observed that disruption upstream of the *uppS-cdsA* operon is significant in both suppressor conditions (Fig. 9B). It was unclear from the insertion pattern the effect that disruption at this locus would have. However, these data suggest that deregulation of the native level of expression of *uppS-cdsA* is restorative in a *yhcB* mutant background.

**Figure 9.**
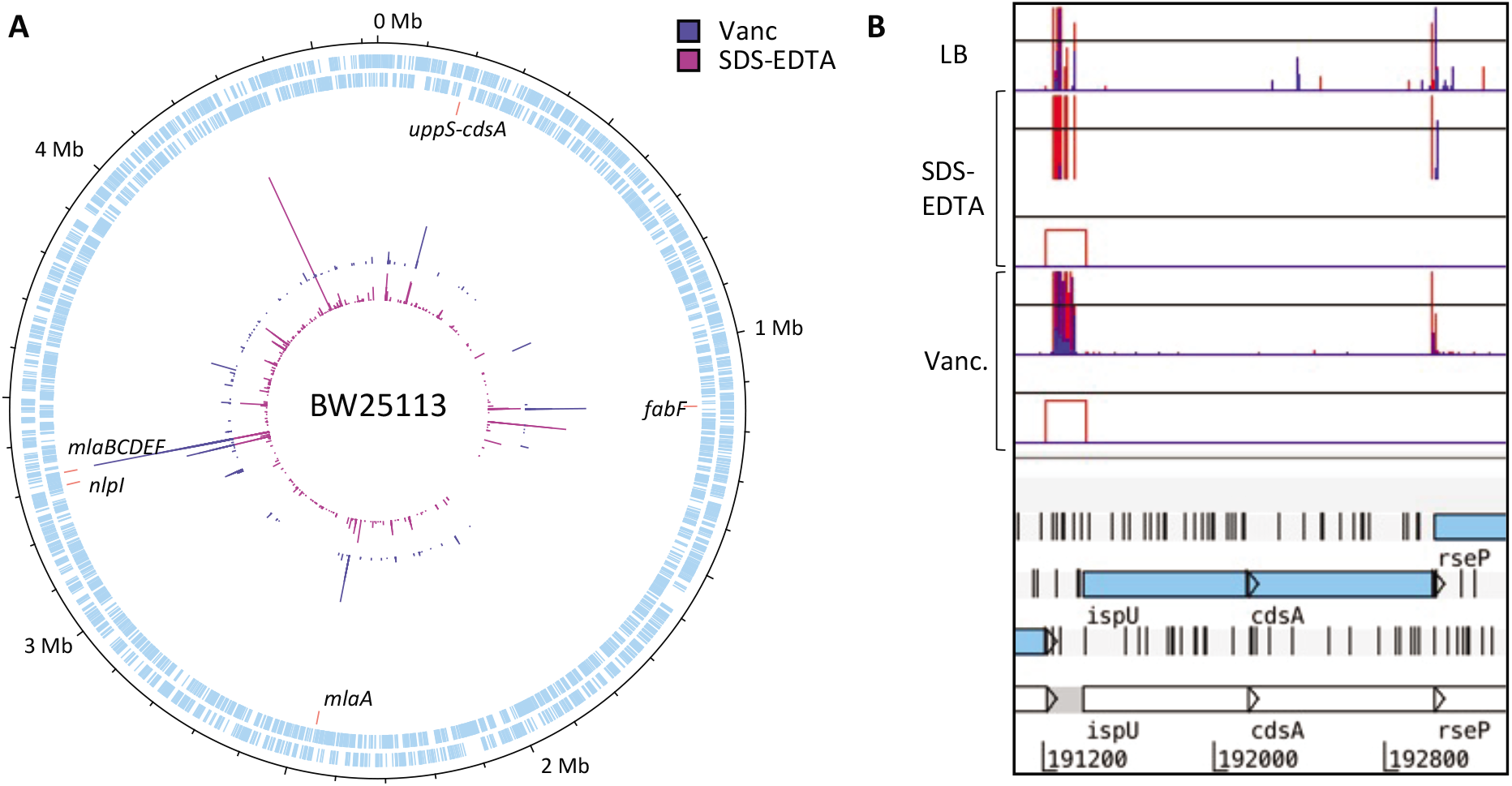
The *uppS-cdsA* operon. (A) Genome map showing the location and frequency of transposon insertion events that restore the viability of a *yhcB* mutant under toxic growth conditions: LB supplemented with vancomycin, or SDS and EDTA. (B) TIS data of the *uppS* (*ispU*) and *cdsA* operon. (Note that *uppS* is annotated as *ispU* in the BW25113 reference genome). Red and blue data correspond with the transposon orientation at the point of insertion. The height of the bar corresponds with mapped sequencing read frequency, capped at 20 in these images. The red boxes underneath each suppressor dataset represent significant differential abundance of insertions of the condition compared to the control, identified by AlbaTraDIS.

Finally, during screening of the transposon mutant library on supplemented agar plates, we also isolated six colonies of the parent Δ*yhcB* strain growing on the control plates of LB supplemented with vancomycin, assumed to be natural revertant suppressors of Δ*yhcB*. The revertant suppression was confirmed by plating on both vancomycin and SDS-EDTA, as well as confirming gene-deletion by PCR of the *yhcB* locus (Fig. S9A-B). We sequenced the genomes of these isolates and identified the mutations listed in Table S8. Two isolates contained either the reported *mlaA*\* mutation or a variation of this mutation (mut4 and mut6), while one isolate had a genome inversion that resulted in separation of the *mla* operon from its promoter region (mut2). The *mla* inversion mutation restored resistance to both vancomycin and SDS-EDTA, consistent with the earlier TIS data, but did not fully restore cell size (Fig. S9A-C). The *mlaA*\* mutation is well characterised and is toxic in a WT background as it results in LPS overproduction as a consequence of increased LpxC stability (May and Silhavy 2018). However, these mutations are not toxic in a Δ*yhcB* mutant and restore cell size and resistance to vancomycin (Fig. S9A-C). We also identified a single nucleotide polymorphism (SNP) in *lpxC* that restored resistance to vancomycin and SDS-EDTA but did not restore cell size (mut1). These data are suggestive of either a defect in LPS synthesis or export, or an imbalance in LPS to phospholipid at the outer membrane of a Δ*yhcB* mutant, rendering the *mlaA*\* mutation non-lethal in this background.

Finally, two isolates each had a SNP that resulted in a single amino acid substitution in CdsA. We used Phyre2 to predict the structure of CdsA (Kelley et al. 2015). The predicted structure was very similar to the experimentally determined structure of CdsA, derived from *Thermotaga maritama*, with an RMSD value of 0.404 A (Fig. S9D) (Liu et al. 2014). The active site of CdsA is a conserved, negatively charged pocket occupied by two cation cofactors that are critical for function (Fig. S9D). Mutations within this region that hinder cation binding result in decreased enzyme activity (Liu et al. 2014). The mutations identified within CdsA in our revertant suppressor mutants (S231L and S239G, respectively) each result in substitution of a serine residue proximal to the metal ion binding site (Fig. S9D). A mutation of S223C in *T. maritama* CdsA, which corresponds with residue S239 in *E. coli*, results in severely reduced CdsA function (Liu et al. 2014). The two mutations within CdsA in the revertant suppressor mutants likely result in decreased function of CdsA. Both mutants with a SNP in CdsA (mut3 and mut5) restored resistance to both vancomycin and SDS and EDTA, in addition to cell size (Fig. S10A-C).

### *Characterisation of a* yhcB-*mutant envelope*

Our genetic screening revealed a connection with phospholipid biosynthesis (*fabF, cdsA*) and trafficking (*mla*), Previous research has shown that fatty acid availability determines the cell size via phospholipid biosynthesis (Vadia et al. 2017). Increased phospholipid production resulted in a larger cell size that was reminiscent of a Δ*yhcB* mutant (Vadia et al. 2017). As such, we hypothesised that phospholipid production might be increased in a *yhcB* mutant. We first investigated whether membrane “lipid ruffles” resulting from increased phospholipid synthesis, and resembling those reported by Vadia et al. (2017), could be detected in a *yhcB* mutant. Following 3 hours of growth in LB, *E. coli* BW25113 and an isogenic *yhcB* mutant were stained with MitoTracker to label internal lipid structures (Vadia et al. 2017). Microscopic evaluation of these cells revealed that *E. coli* BW25113 membranes stained uniformly. In contrast, intensely stained spots indicative of accumulation of lipid structures were observed in the *yhcB* mutant (Fig. S12A). Subsequent imaging of cells by transmission electron microscopy confirmed the aberrant size of *yhcB* mutant cells and revealed the presence of internal membrane structures (Fig. 10A, Fig. S10). These structures were further examined in fast frozen cells after freeze substitution (Walser et al, 2012) avoiding artefacts associated with chemical fixation. Internal membranes showed a range of different morphologies including tubular and vacuolar structures (Fig. 10B, Fig. S11) as well as striking tightly-stacked membrane arrays (yellow arrowheads Fig. S11).

**Figure 10.**
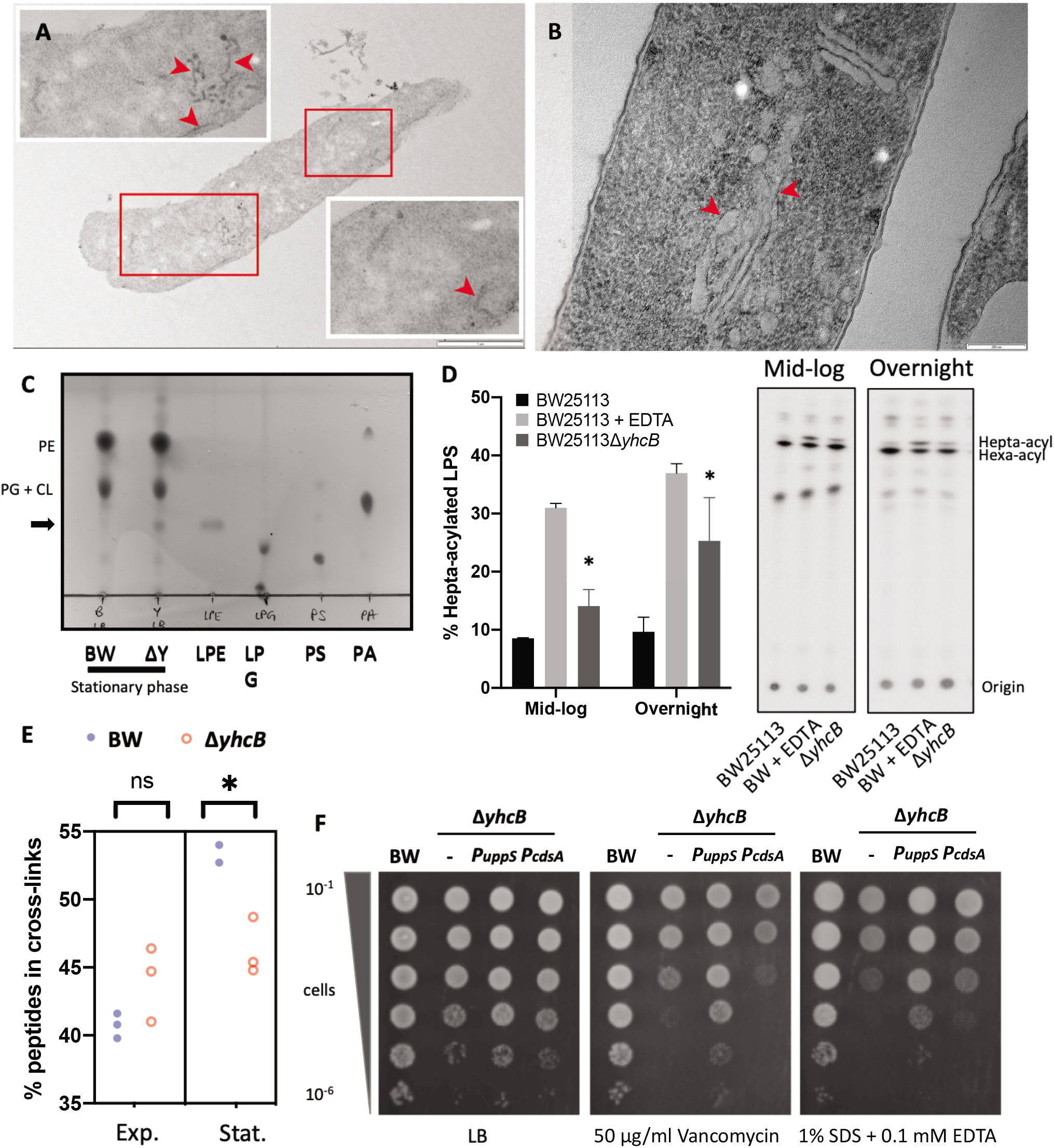
The *yhcB* mutant contains extra membrane and has an altered cell envelope. (A) Transmission electron micrograph (TEM) of a Δ*yhcB* mutant cell with membrane ruffles indicated by red arrows. Scale bar = 1 μm. Additional whole cell images are in Fig. S10. (B) Representative TEM image of a Δ*yhcB* mutant cell processed without primary fixation by fast freezing and freeze substitution. Internal membrane structures indicated by red arrows. Scale bar = 200nm. (C) Phospholipid species extracted from BW25113 (BW) and the Δ*yhcB* mutant (ΔY) grown in LB broth until stationary phase and separated by thin layer chromatography (TLC) using a chloroform:methanol:water (65:25:4) solvent system. Standards of lysophosphatidylethanolamine (LPE), lysophosphatidylglycerol (LPG), phosphatidylserine (PS) and phosphatidic acid (PA) were loaded alongside for comparison. The additional spot in the ΔY sample is indicated with an arrow. (D) TLC/autoradiographic analysis of [^32^P]-labeled lipid A extracted from mid-logarithmic growth and overnight growth cultures of BW25113 and Δ*yhcB* strains grown in LB. As a positive control for lipid A palmitoylation, BW25113 cells were treated with 25 mM EDTA for 10 min prior to extraction. The mean percentage of hepta-acylated lipid A and standard deviation were calculated from samples prepared in triplicate. Student’s *t*-tests: *P<0.05 compared with BW25113. (E) Comparison of the percentage of peptides in peptidoglycan crosslinks between strains at two stages of growth. Unpaired t-test with Welch’s correction: \**P* < 0.05. Exponential phase cells were collected at an OD of 0.4, Stationary phase cells were harvested after 16 h at an OD of ~4.0. (F) Day cultures of BW25113; Δ*yhcB*; Δ*yhcB PuppS* and Δ*yhcB PcdsA* strains grown in LB for 5 h, normalised to an OD_600_ of 1.00 and 10-fold serially diluted before inoculating LB agar plates supplemented with vancomycin or SDS and EDTA. Abbreviations: Phosphatidylethanolamine (PE); Phosphatidylglycerol (PG); Cardiolipin (CL); Exponential phase (Exp.); Stationary phase (Stat.).

Given the increase in phospholipid biogenesis noted above we next investigated the composition of phospholipids species. We extracted the total phospholipid species using the Bligh and Dyer method (Bligh and Dyer 1959), and separated these by thin layer chromatography (Matsumoto et al. 1998). The phospholipid composition of *E. coli* is approximately 75% phosphatidyl ethanolamine, 20% phosphatidyl glycerol and 5% cardiolipin with lysophospholipids typically making up <1% of total phospholipid species (Rowlett et al. 2017; Zheng et al. 2017). The relative ratios of phosphatidylethanolamine, phosphatidylglycerol and cardiolipin were similar in the parent and Δ*yhcB* strains. However, during growth in LB, we observed an additional phospholipid species that was present in both strains, but only the Δ*yhcB* mutant strain in stationary phase (Fig.10B and S10B). This was not observed when the mutant was grown in M9-glucose (Fig. S10C). By using known lipid standards, the additional spot in the Δ*yhcB* mutant stationary phase was identified as most likely lysophosphatidylethanolamine (LPE; Fig. 10B).

LPE can be derived from PE hydrolysis or lipoprotein maturation. However, in a cell with a stressed envelope, the primary source of LPE accumulation occurs via PE hydrolysis (Zheng et al. 2017). Two OM proteins, PldA and PagP, maintain the OM asymmetry by cleaving phospholipids that accumulate in the outer leaflet of the OM. PldA degrades phospholipids while PagP, a Lipid A palmitoyltransferase, cleaves an acyl chain from PE and catalyses its transfer to Lipid A resulting in hepta-acylated LPS (Bishop et al. 2000). Both reactions result in an increase in free LPE. As PagP is usually inactive in the OM and activity is stimulated in response to migration of phospholipids into the outer leaflet (Jia et al. 2004), measuring the amount of hepta-acylated LPS is an established proxy for detecting loss of OM asymmetry (Chong et al. 2015). We quantified the amount of hepta-acylated lipid A relative to hexa-acylated lipid A and identified a significant increase in hepta-acylated lipid A in the Δ*yhcB* mutant compared to the WT; a phenotype that is exacerbated by prolonged growth (Fig. 10C). Together these data suggest that excess phospholipid production in a *yhcB* mutant background results in an increase in cell size and loss of OM asymmetry, resulting in hepta-acylation of lipid A and accumulation of LPE.

Cell membrane synthesis must be coordinated with peptidoglycan biogenesis to maintain cell envelope integrity. As the peptidoglycan layer defines cellular morphology, and we had observed a swollen morphology of *yhcB* mutants, we hypothesised that the peptidoglycan layer of a *yhcB* mutant may be compromised. Indeed, our genetic evidence supports the hypothesis that mutations (*dapF, dacA*) that weaken integrity of the peptidoglycan are lethal. We therefore analysed the muropeptide composition of the cell wall. Cells were grown in LB and collected at both exponential and stationary phase. Peptidoglycan was purified and digested by the muramidase cellosyl, and the resulting muropeptides were analysed by HPLC (Glauner 1988). There were no significant differences in the muropeptide species identified (Fig S11; Table S5). However, we did identify differences in the amount of crosslinking between strains and growth phases (Fig. 10D). While the degree of peptidoglycan crosslinking in the WT strain was increased in stationary phase compared to exponential phase, consistent with the literature (Templin et al. 1999), this same transition was not observed in the *yhcB* mutant, suggesting either a limitation in the rate of peptidoglycan crosslinking or an increased rate of crosslinking hydrolysis in a *yhcB* mutant. The muropeptide analysis, together with the phospholipid analysis, support our finding that a *yhcB* mutant is unable to sustain rapid exponential growth and indicate a defect in the ability to transition into stationary phase.

In addition to a synthetically lethal relationship with genes involved in peptidoglycan assembly, recycling and remodelling (*mepS, mepM, ldcA, nlpl*) our TIS data also indicated that mutations in the ECA biosynthetic pathway that sequester undecaprenol phosphate (Und-P) are lethal in a *yhcB* mutant. We hypothesised that the availability of Und-P might be limited in a Δ*yhcB* mutant, affecting peptidoglycan synthesis (Danese *et al*. 1998; Ramos-Morales *et al*. 2003; Jorgenson *et al*. 2016). In addition to peptidoglycan and ECA biosynthesis, Und-P is also a lipid carrier for O-antigen, however, most *E. coli* K-12 strains, including BW25113, do not have O-antigen as a result of an insertion element in *wbbL* (Liu and Reeves 1994). Therefore, to investigate whether sequestration of Und-P is toxic to a *yhcB* mutant, we tried to restore O-antigen. The transformation efficiency of the *yhcB* mutant with *wbbL* to restore O-antigen was strikingly reduced, suggesting that Und-P limitation is lethal to a *yhcB* mutant (Fig. S10D). One mechanism to suppress Und-P limitation is to upregulate expression of *uppS*, the gene required for Und-P synthesis. Indeed, our TIS data reveal insertions that suppress the *yhcB* phenotype in the promoter region to *uppS*. Furthermore, complementation of *a yhcB* mutant with ectopically expressed *uppS*, but not *cdsA*, suppressed both the vancomycin and SDS-EDTA sensitivity defect of a *yhcB* mutant (Fig. 10E).

During our analysis we noted that *uppS* and *cdsA* are highly co-conserved across the bacterial kingdom (Fig. S12) and share a conserved operon structure in many bacterial phyla (Fig.S13) (Saha et al. 2020). We hypothesise that the upregulation of *uppS* would result in increased expression of *cdsA*, and in turn give rise to excess phospholipid synthesis. This hypothesis is supported by our finding that loss of function mutations in the metal-binding pocket of CdsA restore all *yhcB* defective phenotypes (Fig. 9C).

## Discussion

Here we have applied a high-density mutagenesis screen to comprehensively map the genetic interactions of a single gene, *yhcB*. With thousands of deleterious and restorative genetic interactions discovered, to the best of our knowledge no other gene has been scrutinised genetically at such scale and resolution. Our data reveal that a Δ*yhcB* mutant has a cell envelope permeability defect, dysregulated cell size, and epistatic interactions with multiple pathways of cell envelope biogenesis. The defect introduced by deletion of *yhcB* is most severe during rapid growth in rich media, when the demand for synthesis of cell envelope material is greatest. In addition, toxicity progresses with sustained growth in rich medium, with a decrease in cell viability observed upon transition to stationary phase, a period of growth when *de novo* synthesis machineries are switched off. Taken together, these observations indicate that a *yhcB* mutant is either limited in the ability to recycle existing components of the envelope during synthesis, or limited in *de novo* synthesis of one or more cell envelope components. This hypothesis is supported by the cell growth model put forward by Harris and Theriot (2016) that when cell envelope synthesis is impeded, an increase in cell width is observed decreasing the surface area to cell volume ratio and enabling the same amount of cell volume to be encapsulated by less material.

The pathways for synthesis of the different components of the Gram-negative cell envelope are intrinsically linked: they share several common precursors and tightly interconnected feedback mechanisms (Fig. 11A). If the rate of peptidoglycan biosynthesis is decreased, the rate of LPS biosynthesis needs to be decreased accordingly to avoid the toxic effects of LPS accumulation (Sullivan and Donachie 1984; May and Silhavy 2018). Moreover, a recent paper identified that deletion of the *mla* pathway confers a fitness advantage in cells exposed to fosfomcin, a MurA inhibitor, connecting decreased PG synthesis with a requirement for decreased PL trafficking (Turner et al. 2020). Similarly, when LptC, a component of the LPS export system, is depleted the relative abundance of proteins involved in phospholipid biosynthesis and trafficking are also depleted indicating a feedback mechanism that couples phospholipid synthesis and export with LPS demand (Martorana et al. 2014). In a *yhcB* mutant we observed weakened peptidoglycan integrity and a loss of OM asymmetry, coupled with an increase in cell size and internal phospholipid membrane structures (Fig. 11B). These findings connect YhcB with every layer of the cell envelope.

**Figure 11.**
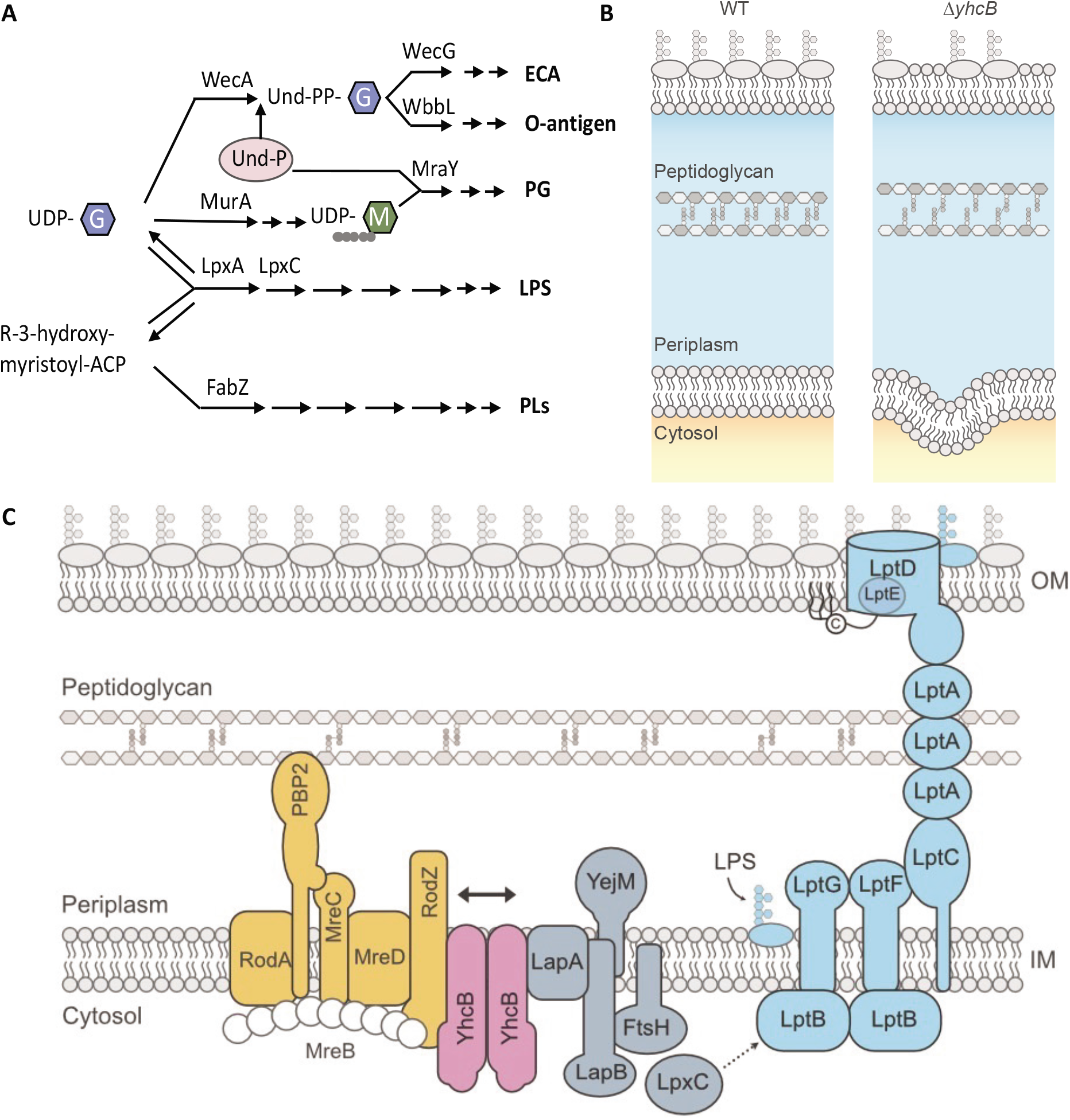
The pathways of cell envelope biosynthesis and relationship with YhcB. (A) Schematic overview of the relationships between the biosynthetic pathways of individual components of the cell envelope. (B) A schematic of the imbalanced cell envelope of a *yhcB* mutant in comparison to wildtype (WT). The *yhcB* mutant suffers from defects in OM asymmetry, excessive phospholipid synthesis and altered crosslinking in the peptidoglycan layer. (C) The positional context of YhcB in conjunction with the reported protein interaction partners of YhcB *in situ*. YhcB interacts with both the elongasome (depicted here as mediated via RodZ, but YhcB is reported to interact with RodA, MreD and MreC in addition to RodZ) and LapA from the complex that regulates LpxC degradation. YhcB is spatially positioned to coordinate peptidoglycan expansion with LPS export. Abbreviations: Enterobacterial Common Antigen (ECA); Lipopolysaccharide (LPS); Peptidoglycan (PG); Phospholipid (PL); Undecaprenyl phosphate (Und-P); Undecaprenyl diphosphate (Und-PP); Uridine diphosphate (UDP); N-acetylglucosamine (G); N-acetylmuramic acid (M); Acyl Carrier Protein (ACP).

Our observations of excess phospholipid membranes and an increased cell size in a *yhcB* mutant resemble the phenotype reported by Vadia *et al*. (2017) that results from increased phospholipid synthesis. We hypothesise that the increase in phospholipid production results in loss of OM asymmetry, with substantial amounts of phospholipid integrating into the outer leaflet of the OM to compromise the barrier function of the cell envelope. This hypothesis is supported by several lines of evidence. First inhibition of phospholipid synthesis by mutations within the metal ion binding site of CdsA, or loss of FabF, which regulates membrane fluidity, are restorative to a *yhcB* mutant. Furthermore, treatment with sub-inhibitory concentrations of cerulenin, an antibiotic that targets the FabB and FabF elongation step of fatty acid synthesis, is beneficial to growth of a *yhcB* mutant (Nichols et al. 2011). Second, a *mlaA*\* mutation, which increases mislocalisation of phospholipids at the OM triggering subsequent phospholipid degradation by PldA and is toxic in a wild-type background, is tolerated in a *yhcB* mutant, restores cell size, and re-establishes the barrier function of the OM. Third, an increase in LPS production through the loss of the essential protease FtsH, the introduction of *mlaA*\* which upregulates LpxC production, or by mutation of its target LpxC can be tolerated. Overall, mutations that result in a decrease of phospholipid biosynthesis and trafficking, or an increase in LPS biosynthesis are restorative, indicative of an underlying imbalance in PL to LPS levels.

We found that mutations that weaken PG integrity are lethal in a *yhcB* mutant indicating that the rate of PG synthesis is also not coordinated with cell expansion. While the composition of the PG in a *yhcB* mutant resembles that of the WT, our data reveal that loss of PG synthesis by the elongasome is restorative. In contrast, loss of PG synthesis by the divisome had a negative effect while mutations that stimulate PG synthesis via the divisome are restorative. Given the location of YhcB and the phenotypes of a deletion mutant, an attractive hypothesis is that YhcB is involved in switching PG synthesis between the elongasome and divisome. In support of this hypothesis a recent paper reported the interaction and co-purification of YhcB with proteins of the divisome (Mehla et al. 2021).

Another attractive hypothesis is that YhcB has a role in sensing the availability of Und-P, or recycling/synthesis of Und-P. This latter hypothesis is supported by several observations. First, mutations in the ECA biosynthesis pathway that sequester Und-P and limit the available pool are lethal. Second, restoration of WbbL activity is toxic in a *yhcB* mutant, suggesting that Und-P availability is limited. Third, a *yhcB* mutant is reported to be more sensitive to bacitracin (Nichols et al. 2011), which targets BacA. BacA is responsible for recycling approximately three quarters of Und-PP back to Und-P following release of its substrate in the periplasm. Fourth, deletion of *mrcB* in cells with low levels of Und-P results in heterogenous cell lysis, and *mrcB* was identified in the list genes in which an epistatic interaction with *yhcB* causes a fitness defect. These data are consistent with the finding that only a sub-population of the Und-P depleted *mrcB* mutants lyse (Jorgenson et al. 2019). Lastly, upregulation of *uppS*, but not *cdsA*, was universally restorative in a *yhcB* mutant.

The observation that the *uppS-cdsA* operon structure is conserved across bacteria reveals the tight linkage between phospholipid production and the synthesis of polysaccharide components of the cell envelope. The observation that cell size, resistance to vancomycin, and resistance to SDS-EDTA can be decoupled in suppressor screens indicate that YhcB interacts with multiple cell envelope biogenesis pathways. Indeed, in addition to forming a dimer and reports of larger homo-multimer structures (Maddalo et al. 2011; Li et al. 2012; Mehla et al. 2021), YhcB is known to interact with several components of the elongasome (Li et al. 2012), and has also been identified as an interaction partner of LapA (Hu et al. 2009; Li et al. 2012). LapA (YciS) forms a complex with LapB (YciM), FtsH and YejM to regulate LpxC stability and thus LPS abundance (Clairfeuille et al. 2020; Fivenson and Bernhardt 2020; Guest et al. 2020). The role of LapA is unclear but deletion of *lapA* results in minor accumulation of incomplete LPS precursors (Klein et al. 2014). The interaction of YhcB with the elongasome and the LpxC regulation complex via LapA positions YhcB at the interface between two of the major complexes that coordinate envelope synthesis (Fig. 11C). This hypothesis is supported by the observation that deletion of different YhcB domains gives rise to different chemical sensitivity profiles.

In conclusion, YhcB plays an important regulatory function at the interface between all of the cell envelope biogenesis pathways, and this function is mediated by physical interactions between members of the elongasome and LPS regulation systems. While the precise molecular events facilitating the regulation of these pathways has yet to be elucidated, as *yhcB* is conserved among Gammaproteobacteria, is required for tolerance to polymyxin B, colistin, and vancomycin, and is necessary for the colonisation of different hosts (Harvey et al. 2011; Brooks et al. 2014; Jana et al. 2017), it provides a novel antimicrobial target for exploitation against clinically important pathogens.

## Materials and Methods

### Strains, media and growth conditions

The parent strain for this work was *E. coli* K-12 strain BW25113. Gene deletion mutants were constructed by P1 transduction and using the Keio library strains as donors (Baba et al. 2006; Thomason et al. 2007). The kanamycin cassette was removed using the pCP20 vector (Datsenko and Wanner 2000). The mutant was confirmed by PCR and Sanger sequencing. Bacteria were grown in Luria-Bertani (LB: 10 g tryptone, 5 g yeast extract, 10 g NaCI) medium or on LB plates (LB supplemented with 1.5% nutrient agar) and incubated at 37°C. When required, media were supplemented with 50 μg/ml kanamycin or 100 μg/ml carbenicillin. The M9-glucose recipe used was: 1x M9 salts (Sigma Aldrich), 200 μl filter sterilised 1 M MgSO_4_, 10 μl filter sterilised 1 M CaCl_2_, and 2 ml of either a 20% (w/v) D-glucose or 20% (v/v) glycerol solution per 100 ml. For micro-dilution spot plates, unless otherwise stated, bacteria were grown overnight in 5 ml LB medium at 37°C with aeration. Cultures were normalised by optical density to an OD_600_ of 1.00, 10-fold serially diluted in LB, and 2 μl of each dilution was inoculated onto LB agar plates.

### Complementation assays

The BW25113 and *yhcB* strains were transformed with a pBAD-Myc-His-A plasmid with and without the *yhcB* coding sequence under the control of the arabinose promoter. O-antigen was restored as described previously (Browning et al. 2013). In brief, cells were transformed with a pET20b plasmid DNA encoding *wbbL* Additional copies of *cdsA* and *uppS were* introduced, independently, on a pASK-IBA2C plasmid under the control of a tetracyclin-inducible promoter.

### Polymyxin B TIS screen

An amended version of the Andrews broth microdilution protocol in 96-well plate format, adjusting the starting inoculum to an initial OD_600_ of 0.05 and the growth medium to LB, was used to identify an initial inhibitory concentration range or polymyxin B (Andrews 2001). Growth curve experiments of *E. coli* BW25113 were then repeated in 50 ml LB supplemented with polymyxin B in glass flasks, reflecting the conditions of the TIS screen. A concentration of 0.2 μg/ml polymyxin B was identified as sub-inhibitory for growht of BW25113. The transposon library was inoculated into 50 ml LB medium with and without 0.2 μg/ml polymyxin B (Sigma Aldrich), in duplicate, at a starting optical density of 0.05, equivalent to a copy number of ~2,500 of each mutant, and grown to OD_600_ = 1.00, at 37°C with aeration. Cells were harvested, and genomic DNA prepared for sequencing. As we knew the density of this library (Goodall et al. 2018), we estimated the number of mapped reads, and therefore sequencing coverage, needed to ensure sufficient sampling of the library using the equation 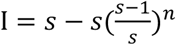, where I = insertions, and n = number of mapped reads (Fig. S1B). For a theoretical library of 1 million mutants, assuming no loss of mutants and an equal chance of each transposon junction being sampled, ~2.3M reads are needed to ensure sampling of 90% of the library, with diminishing returns with further sequencing. Obtaining >4.2 M reads should enable sampling of 99% of the possible unique insertion sites. Therefore, we collected >2 M sequencing reads per replicate, and a combined total of >4.2 M reads per condition, to give us confidence that any observed loss of mutants is not due to insufficient sampling.

### Microscopy

Samples were taken directly from overnight cultures grown at 37°C in their respective media for 16 h and diluted to an OD_600_ of 0.10. 5 μl cells were spread on a 1 mm glass slide pre-treated with 5 μl poly-L-lysine (Sigma Aldrich). A Nikon 9Oi eclipse microscope was used to capture differential interference contrast (DIC) images of cells, using a 40x objective lens with a Nikon immersion oil. These images were collected at the University of Birmingham. Cell dimensions were measured using ImageJ (Schindelin et al. 2012). Fluorescent labelling of lipids was previously described by Vadia *et al*. (2017). These experiments were done at the University of Queensland. M9-glucose overnight cultures were diluted into fresh LB medium and after 2 h of growth at 37°C, 0.2 μM MitoTracker® Green FM (Invitrogen) was added to each sample to label lipids. After incubation for a further 1 h, bacteria from 1 ml samples were harvested by centrifugation and resuspended in 500 μL of FM™ 4-64FX (Invitrogen) at 5 μg/mL, to label *E. coli* membranes, and incubated for 5 min at room temperature. Bacteria were embedded on 1% agarose pads and imaged using a confocal microscope Inverted LSM 880 Fast Airyscan (63x/1.40 OIL). Electron microscopy was performed at the University of Queensland. Conventional fixation and embedding in resin was undertaken using a protocol adapted from (Fassel et al. 1997). Briefly, bacteria were applied to dishes with glass coverslips coated in poly-L-lysine and then fixed in 2% PFA, 2.5% Glutaraldehyde, and 0.075% Ruthenium Red. Samples were then washed 3×10min in PBS and postfixed in 1% osmium, then again in 1% osmium with 3% potassium ferricyanide. Bacteria underwent serial dehydration in increasing concentrations of ethanol and were then infiltrated and embedded with Epon resin. Fast freezing and freeze substitution of bacteria was undertaken as described previously (Ariotti et al. 2012; Walser et al. 2012). Briefly, bacteria were applied to carbon-coated poly-L-lysine treated sapphire discs and frozen in a Leica EMPACT 2 high pressure freezer. They were then freeze substituted in a EM AFS 2 system (Leica Microsystems GmbH, Wetzlar, Germany) and embedded into Epon resin. Ultra-thin sections were obtained using a Leica Ultracut UC6 Ultramicrotome and micrographs acquired on a JEOL 1011 transmission electron microscope equipped with a Morada CCD camera.

### Phospholipid extraction

Total phospholipids were extracted using an amended version of the Bligh-Dyer method (Bligh and Dyer 1959). 10 ml of culture at an OD_600_ ~ 4.00, or 100 ml of culture at an OD_600_ of 0.40 was centrifuged at 4°C to harvest cells. The supernatant was discarded, and pellet resuspended in 1 ml ddH_2_O and transferred to a glass tube. 1.25 ml chloroform and 2.5 ml methanol were added to the sample using glass pipettes. The sample was vortexed for 20 s to create a single-phase solution which was then incubated at 50°C for 30 min. A further 1.25 ml chloroform and 1.25 ml water was added to the sample to create a 2-phase solution and incubated again at 50°C for 30 min. The sample was centrifuged at 400 x *g* for 10 min at RT and the lower organic phase containing phospholipids was transferred to a new glass tube using a glass Pasteur pipette. Chloroform was evaporated by placing the glass tubes in a heat block at 50°C under a stream of nitrogen.

### Thin Layer Chromatography

10 μl of sample was spotted onto the origin of a TLC silica gel membrane (Merck) using a 5 μl glass capillary tube (Sigma Aldrich). Once dry, the membrane was transferred to an equilibrated solvent system of 65:25:4 chloroform:methanol:water (Matsumoto et al. 1998). The samples were separated until the solvent front had migrated sufficiently from the origin then the membrane was removed from the solvent tank and was air dried at RT. Samples were stained with phosphomolybdic acid (PMA) 10% solution in ethanol and heated with a heat gun to activate the PMA until lipid species were visible. Lipid standards were purchased from Avanti Polar Lipids and handled according to manufacturer instructions.

### Lipid A palmitoylation assay

Lipid A labelling, extraction and analysis were done as described and demonstrated previously (Chong et al. 2015). As a positive control, BW25113 was exposed to 25 mM EDTA for 10 min to induce PagP mediated palmitoylation of Lipid A, before harvesting cells by centrifugation.

### Peptidoglycan extraction and analysis

Cell cultures were grown in LB medium at 37°C and harvested at an OD of 0.4 and 4.00, in triplicate. After pelleting the cells and resuspending in ice-cold water, the cell suspension was dropped into 8% boiling SDS solution. Peptidoglycan was purified, and muropeptides were released with cellosyl and analysed by HPLC as described (Glauner 1988).

### Construction of transposon-mutant libraries

10 ml of 2x TY broth was inoculated with a single colony and grown overnight at 37°C with aeration. The 10 ml overnight culture was used to inoculate 800 ml 2x TY broth in a 2 L flask and grown at 37°C with aeration until OD_600_ 0.6-0.9. At the desired OD, cells were collected and stored on ice for 30 min before centrifugation to pellet the cells. Electrocompetent cells were prepared by repeatedly centrifuging cells and resuspending in decreasing amounts of ice-cold 10% (v/v) glycerol. The final resuspension was in 1 ml of 10% glycerol resulting in a dense ~2 ml cell culture. Aliquots of 200 μl cells were distributed between 1.5 ml microcentrifuge tubes. 0.2 μl EZ-Tn5™ transposome (Epibio) was mixed with each aliquot of cells and incubated on ice for 30 min. Samples were transferred to pre-chilled 2 mm gap electroporation cuvettes (Cell Projects Ltd.). Cells were pulsed at 2200 V and 2 ml of pre-warmed SOC medium was immediately added to the sample for recovery. Samples were transferred to a 15 ml falcon tube to allow for maximum aeration and were incubated at 37°C for 2 h. 5 ml of LB broth was added to each 15 ml falcon tube. ~4-5 drops of cells (equivalent to ~200 μl) were spread per LB agar plate supplemented with 50 μg/ml kanamycin. Sufficient cells were inoculated per plate to form non-touching single colonies. Plates were incubated overnight at 37°C for 18 h. Following incubation, 500 μl 30% glycerol-LB broth was added to each plate and using a ‘hockey-stick’ spreader, colonies were scraped off the surface of the agar plate and pooled. Cells were mixed thoroughly before storing at −80°C.

### *Screening of the* yhcB *transposon-mutant library to identify suppressor mutations*

Cell cultures were resuspended in 1 ml of OD_600_ = 1.00, 200 μl of culture was plated across 5x LB agar plates with or without additional chemical stresses. Assuming a library of ~500,000 mutants (as judged from the unique insertions), this number of cells equates to approximately 2,000 independent copies of each individual transposon mutant. We chose a plate-based screening method as this minimises competition between mutants, allowing for identification of slow growing mutants in addition to strong suppressor mutations. Cultures were diluted to such quantities that single colonies of transposon mutants could be isolated on the plates supplemented with stresses to prevent the growth of satellite colonies and therefore false positive results. Plates were incubated at 37°C for 24 h. Colonies were scraped and pooled for sequencing.

### TIS sequencing

Genomic DNA (gDNA) was extracted and quantified using Qubit™ dsDNA HS Assay kit (Invitrogen). 1 μg of gDNA was fragmented by mechanical shearing using a bioruptor (Diagenode) using the shearing profile 30 s ON, 90 s OFF at low intensity, resulting in DNA fragments with an average length of ~300 bp. Fragmented DNA was end-repaired using the NEBNext Ultra I kit (New England Biolabs). The fragments were then ligated with an adapter, and the sample was purified in a size-selection step using AMPure XP SPRI beads (Beckman Coulter) before transposon-junctions were enriched by PCR with primers specific for the transposon and the adapter. The Transposon-gDNA junctions were prepared for sequencing by PCR addition of Illumina adapters using the NEBNext Multiplex Oligos for Illumina (New England Biolabs). However, the forward Universal primer was replaced with custom primers that include the Universal primer sequence followed by a 6-9 nucleotide barcode (to introduce complexity and stagger the start of the transposon sequence) and a 22 nt sequence with homology to the transposon at the 3’ end of the primer. Samples were purified after each PCR step using SPRI beads at a ratio of 0.9:1 beads to sample. Finally, the sample was quantified using the KAPA Library Quant Kit (Illumina) Universal qPCR Mix (Kapa Biosystems). Samples were pooled, denatured and diluted to 18 pM and sequenced using an Illumina MiSeq, with 5% (v/v) 20 pM PhiX (Illumina), using 150 cycle v3 cartridges. Data are available at the European Nucleotide Archive (accession: PRJEB43420).

### TIS analysis

Data were first demultiplexed using the Fastx barcode splitter, to remove the 5’ end barcode (inline index) unique to each sample (Pearson et al. 1997). The BioTraDIS analysis package (version 1.4.5) was used for the remaining data processing (Barquist et al. 2016). We allow for up to 4 bp mismatches in the transposon pattern matching step. When successfully identified transposon sequences have been identified, the transposon is trimmed and the remaining read is mapped to the BW25113 reference genome using bwa (accession CP009273.1) to generate insertion plot files. These data can be viewed online at our browser: http://tradis-vault.qfab.org/. The plot files are input to the tradis_gene_insert_sites script to calculate the number of insertions per gene (insertion index score). Unless otherwise stated, 5% trim was applied to both the 5’ and 3’ end of each gene. The tradis_essentiality.R script was used to calculate the probability of belonging to each mode representing essential and non-essential gene populations respectively. The tradis_comparison.R script was used to compare read depth between control and condition samples, per gene, using a threshold of >2-fold change and a Q-value < 0.01. Plot files generated by the BioTraDIS analysis were also used as inputs for AlbaTraDIS. AlbaTraDIS (version 1.0.1) was used for comparative analysis between suppressor TIS datasets (Page et al. 2020). The reference genome annotation was used to define gene boundaries. We set the minimum log counts per million threshold at 10, for all other settings the default conditions were used.

### Whole genome sequencing

Whole genome sequencing was done by MicrobesNG, University of Birmingham, UK, using Illumina platforms generating short read data. Sequencing data were aligned to the *E. coli* BW25113 reference genome available from the NCBI database (CP009273.1) using bwa mem and then converted to bam files and sorted and indexed using SAMtools (Li et al. 2009; Langmead and Salzberg 2012). Data are available at the European Nucleotide Archive (accession: PRJEB43420). The programs Snippy and VarScan were used to identify SNPs and indels, BreSeq was used to identify large chromosomal rearrangement events (Koboldt *et al*. 2012; Deatherage & Barrick 2014; Seeman 2015).

### Phylogenetic analysis

All searches were performed against the UniProt database of Reference Proteomes (v2020_06, 02-Dec-2020), which includes only complete proteomes of reference organisms, to confidently determine presence as well as absence. Search was performed with PF06295.13 using hmmsearch (3.1b2; (Eddy 2011)) with the --max setting. Search results were analysed manually as well as by all-against-all blast and subsequent clustering in cytoscape, to determine the search cutoff used. Taxonomy information was equally obtained from the UniProt Reference Proteomes ftp site (v2020_06, 02-Dec-2020). Phylogenetic inference was performed on the sequences aligned via mafft (Katoh and Standley 2013), and tree calculation using IQ-TREE (Nguyen et al. 2015) with the built-in model test (Kalyaanamoorthy et al. 2017), which resulted in LG+G4. Tree visualisation was performed using iTol (https://doi.org/10.1093/nar/gkz239).

## Acknowledgements

We thank Dr. Eugenio Sanchez-Moran at the University of Birmingham for use of his microscope and we thank the University of Queensland Microscopy department at the Institute of Molecular Biosciences for assistance in setting up the lipid-labelling microscopy experiments. In particular, thank you Dr. Nicholas Condon for your time and teaching. The authors acknowledge the use of the Microscopy Australia Research Facility at the Centre for Microscopy and Microanalysis at The University of Queensland. In particular, we thank Rick Webb for expert help with cryoEM processing. Light microscopy was performed at the Australian Cancer Research Foundation (ACRF)/Institute for Molecular Bioscience Cancer Biology Imaging Facility, which was established with the support of the ACRF. This research was supported by the Midlands Integrative Biosciences Training Partnership (MIBTIP, BBSRC) Ph.D. program who sponsored E.G. This project received funding from the European Union’s Horizon 2020 research and innovation programme under the Marie Skłodowska-Curie grant agreement No 721484 awarded to WV and IRH. B.Z. was supported by a CSC-UQ PhD Scholarship (China Scholarship Council). This work was also supported by the National Health and Medical Research Council of Australia (grants APP1140064 and APP1150083 and fellowship APP1156489 to R.G.P.). RGP is supported by the Australian Research Council (ARC) Centre of Excellence in Convergent Bio-Nano Science and Technology CE140100036. We thank MicrobesNG for providing a fantastic service generating WGS data, especially Dr. Emily Richardson for your patience and sharing your expertise.

---

**Supplementary Figure 1.**
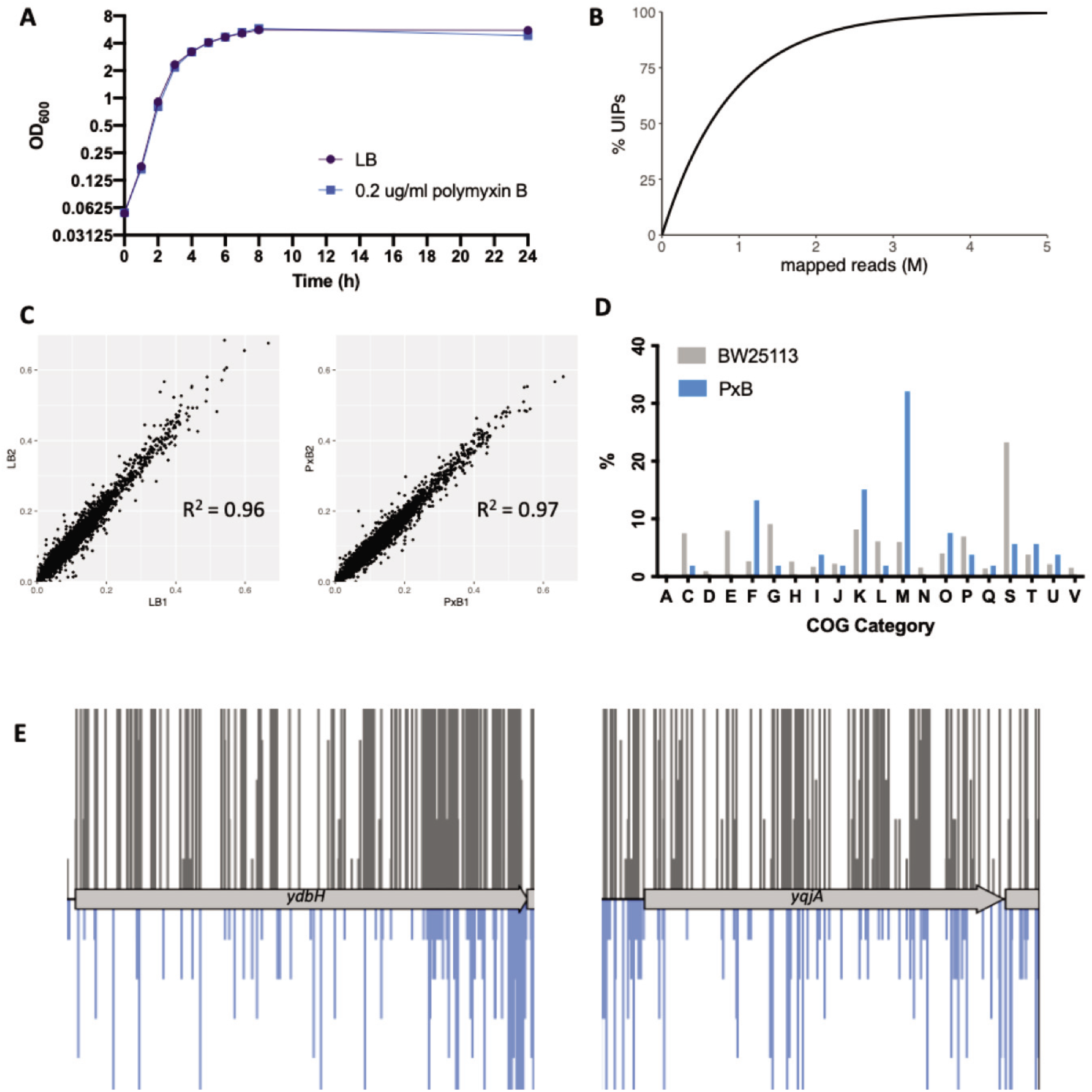
Screening the BW25113 transposon library in sub-inhibitory concentrations of polymyxin B. (A) 0.2 μg/ml of polymyxin B in 50 ml LB does not inhibit growth of *E. coli* BW25113, under the conditions used to screen the library. One representative growth curve is shown, consistent with 5 repeats. (B) Calculation of the number of mapped reads needed to sample a given percentage of the transposon library. The equation 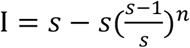 was used to estimate the number of mapped reads required to sample a given proportion of the dataset, s = 1,000,000 was taken as the total number of possible mutants (total transposon insertion sites). I = insertions identified, n = number of mapped reads. This data was used to calculate the approximate percentage of unique insertions identified for a given number of mapped reads, for a library of 1,000,000 unique mutants. (C) Comparison of insertion index scores (a measure of insertion density) per gene for each replicate of the library either exposed to LB only, or LB supplemented with polymyxin B. The scatter plots show the correlation coefficient for insertion density of each gene between replicates. (D) The relative abundance of Cluster of Orthologous Groups (COG) categories for all genes of BW25113 (grey) and for the 54 genes identified as required for growth in sub-inhibitory concentrations of polymyxin B (PxB; blue). (E) The transposon insertion profiles of *yqjA* and *ydbH* following outgrowth in LB only (grey, above) or in LB supplemented with polymyxin B (blue, below). The transposon insertion position along each gene is marked by a vertical line. The vertical line size corresponds with read depth, with visibly fewer transposon-insertion sites identified within *ydbH* and *yqjA* following outgrowth in polymyxin B.

**Supplementary Figure 2.**
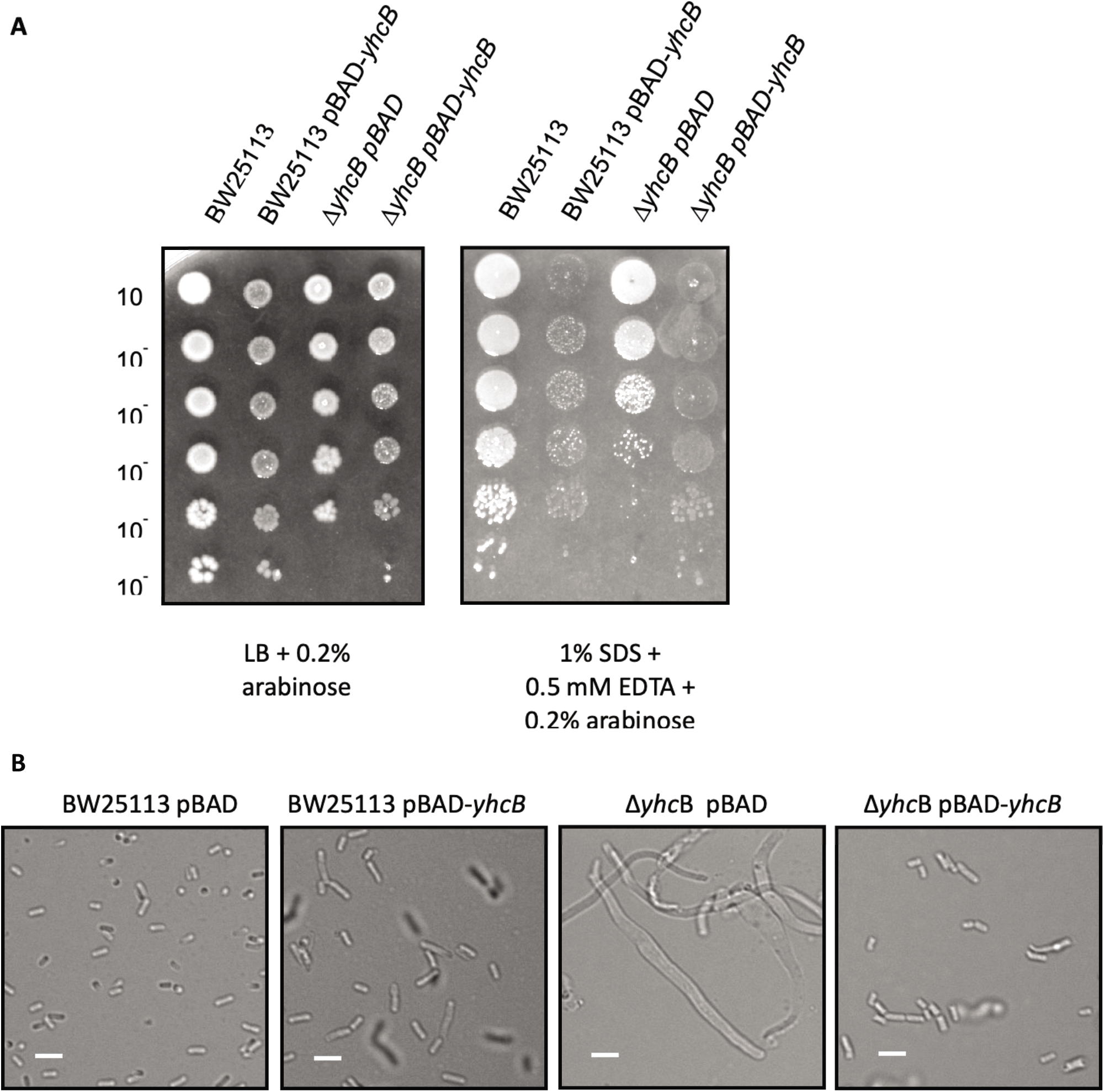
Complementation of a Δ*yhcB* defect. (A) 10-fold serial-dilution of overnight cultures grown in LB, inoculated onto LB agar plates supplemented with 0.2% arabinose, with and without 1% SDS + 0.5 mM EDTA. Strains are carrying a pBad-Myc-His-A with or without the *yhcB* CDS under the control of an arabinose promoter. (B) DIC microscopy images of day cultures grown in LB supplemented with 0.4% arabinose and 100 μg/ml carbenicillin (to maintain plasmids), with a5μm scale bar for reference.

**Supplementary Figure 3.**
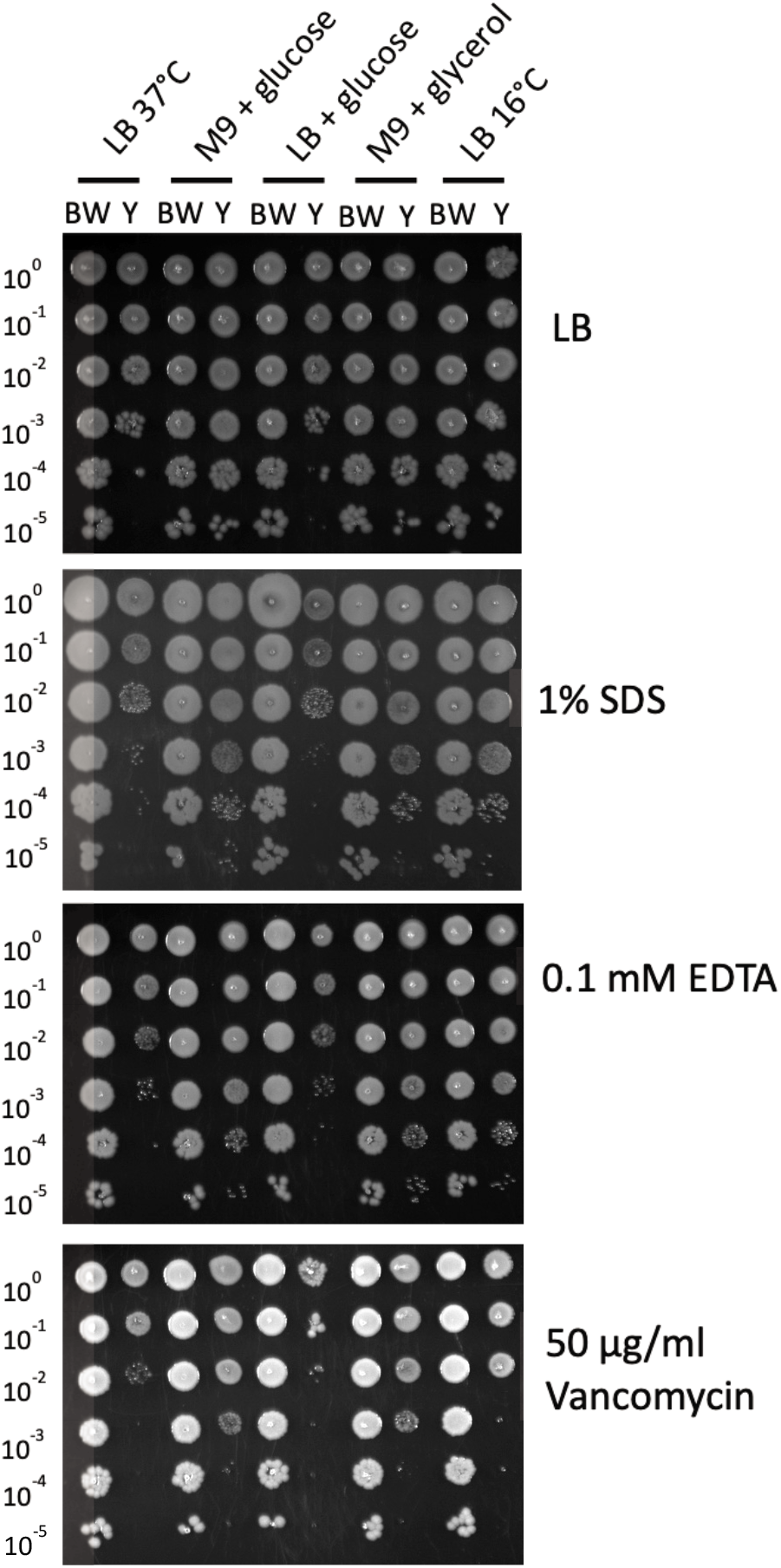
The effect of different growth media on phenotype. 10-fold serial-dilution of overnight cultures of the parent strain *E. coli* BW25113 (BW) and Δ*yhcB* strain (Y) grown to late stationary phase under different conditions (LB at 37°C; M9 + 0.4% glucose at 37°C; LB + 0.4% glucose at 37°C; M9 + 0.4% glycerol at 37°C; LB at 16°C) normalized to an OD_600_ of 1.00 and inoculated onto LB agar plates supplemented with and without 1% SDS; 0.1 mM EDTA or 50 μg/ml vancomycin.

**Supplementary Figure 4.**
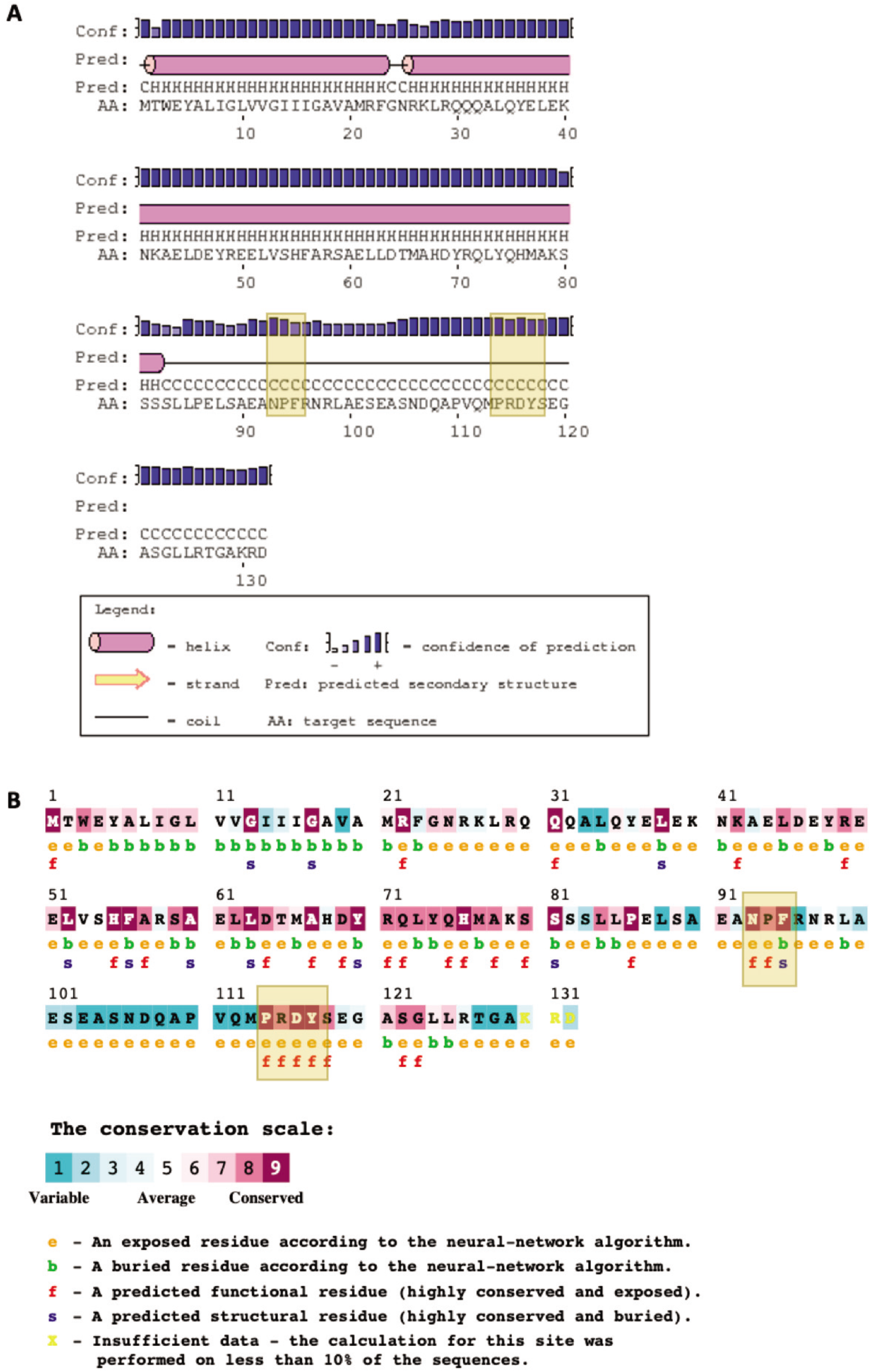
Domains and conserved motifs of YhcB. (A) PSIPRED secondary structure prediction of YhcB (B) Conserved residues of YhcB predicted by ConSurf. Conserved ‘NPF’ and ‘PRDY’ motifs are highlighted with amber boxes in both panels.

**Supplementary Figure 5.**
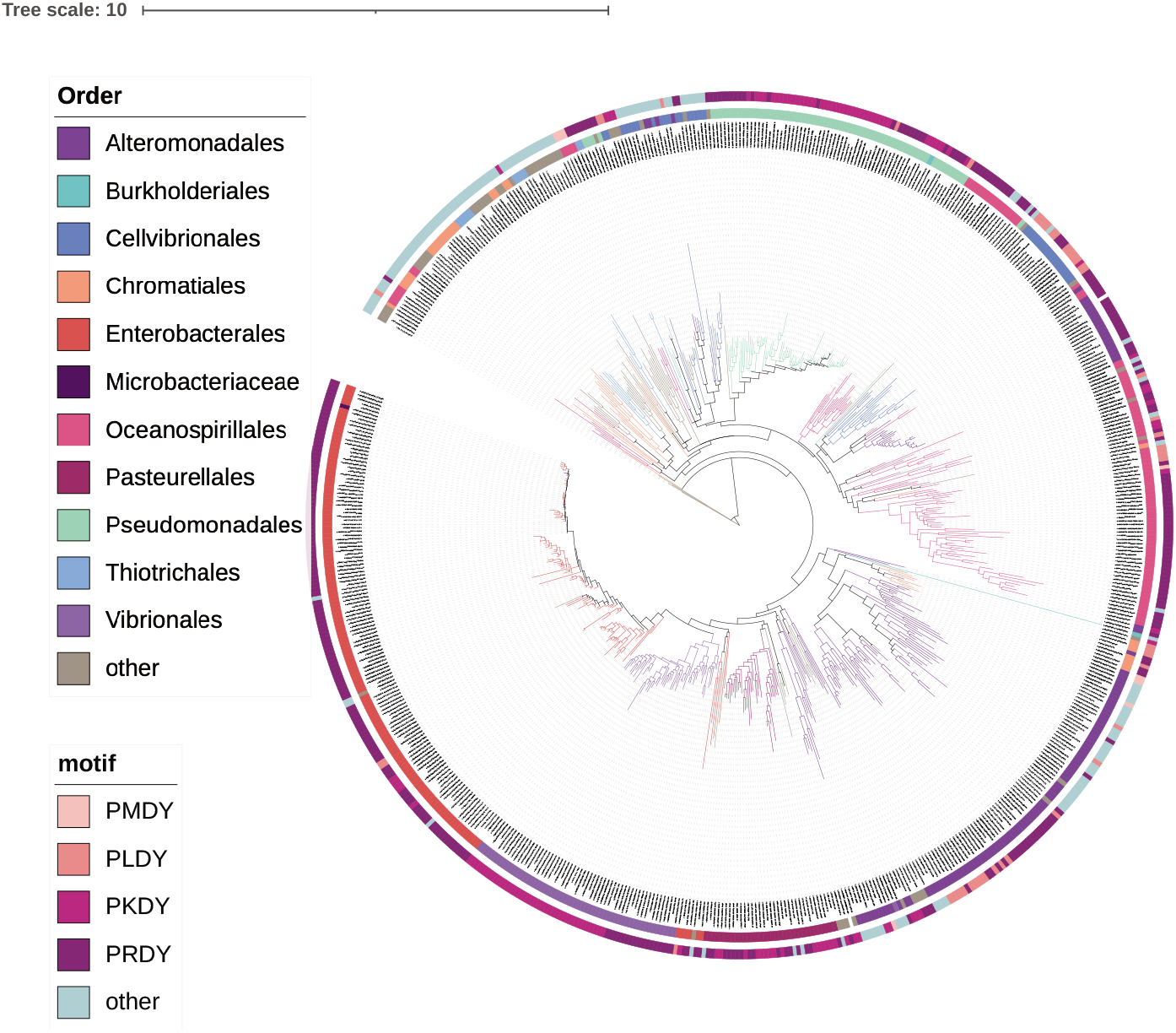
Conservation of the PRDY domain. Phylogenetic analysis displaying conservation of YhcB in bacterial reference genomes. Branches of the tree, and second outermost track, are coloured according to taxonomic Order. The outermost track is coloured according to the amino acid residues within the ‘PXDY’ domain conserved among species. The label ‘other’ represents those that had a different sequence to the four listed.

**Supplementary Figure 6.**
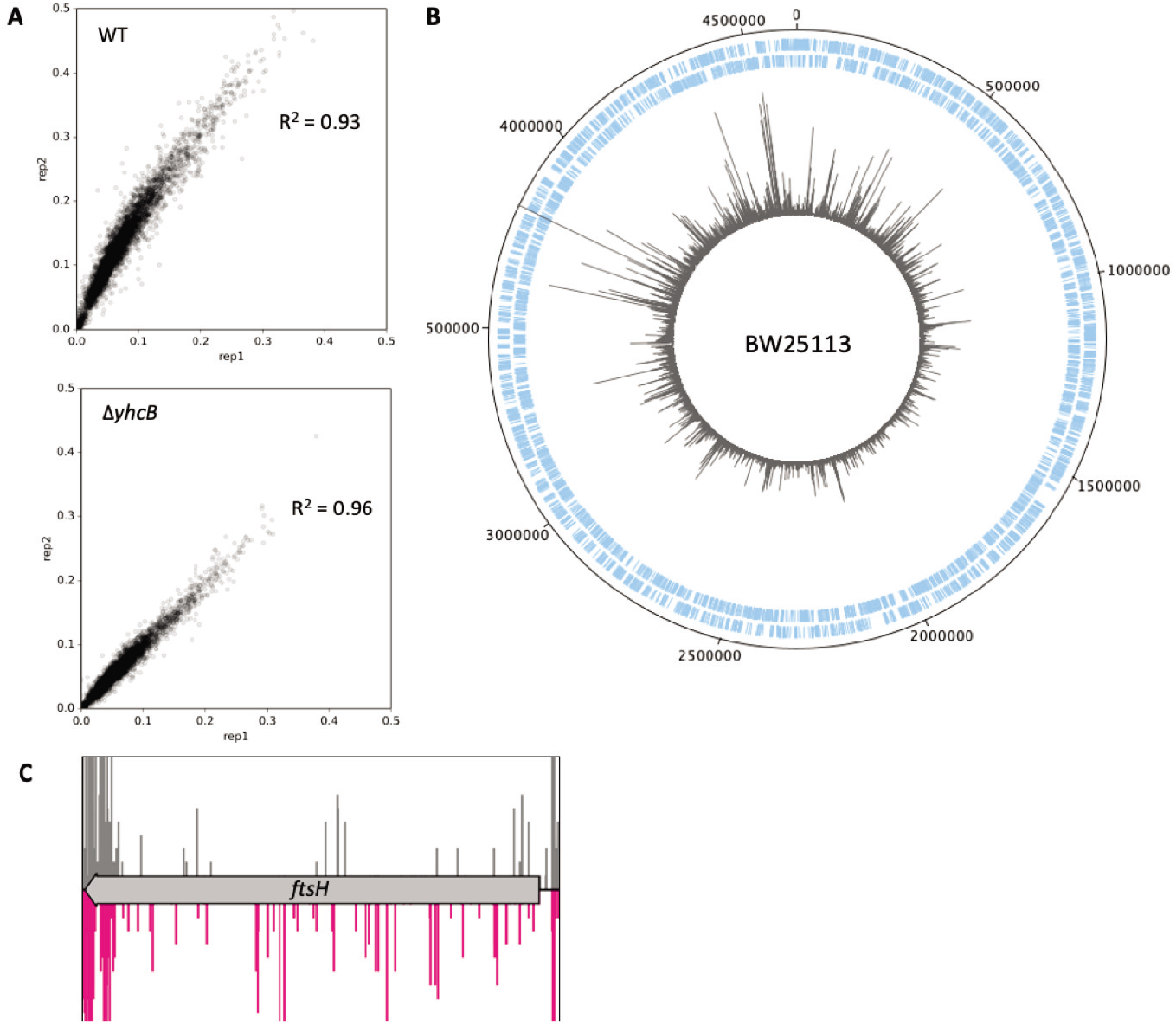
Construction of a transposon library in BW25113 parent strain. (A) Comparison of insertion index scores (IIS) between technical replicates of each library. (B) The BW25113 transposon library. A genome map of BW25113 starting at the annotation origin, with the sense and antisense coding sequences of BW25113 shown in blue, respectively, and the position and frequency of sequenced transposon insertion events shown in grey. (C) The transposon insertion profile of *ftsH* shown for both libraries. Peaks represent the abundance of detected transposon insertion events for each library: WT (grey, above) and *yhcB* (pink, below) with read frequency capped at 10.

**Supplementary Figure 7.**
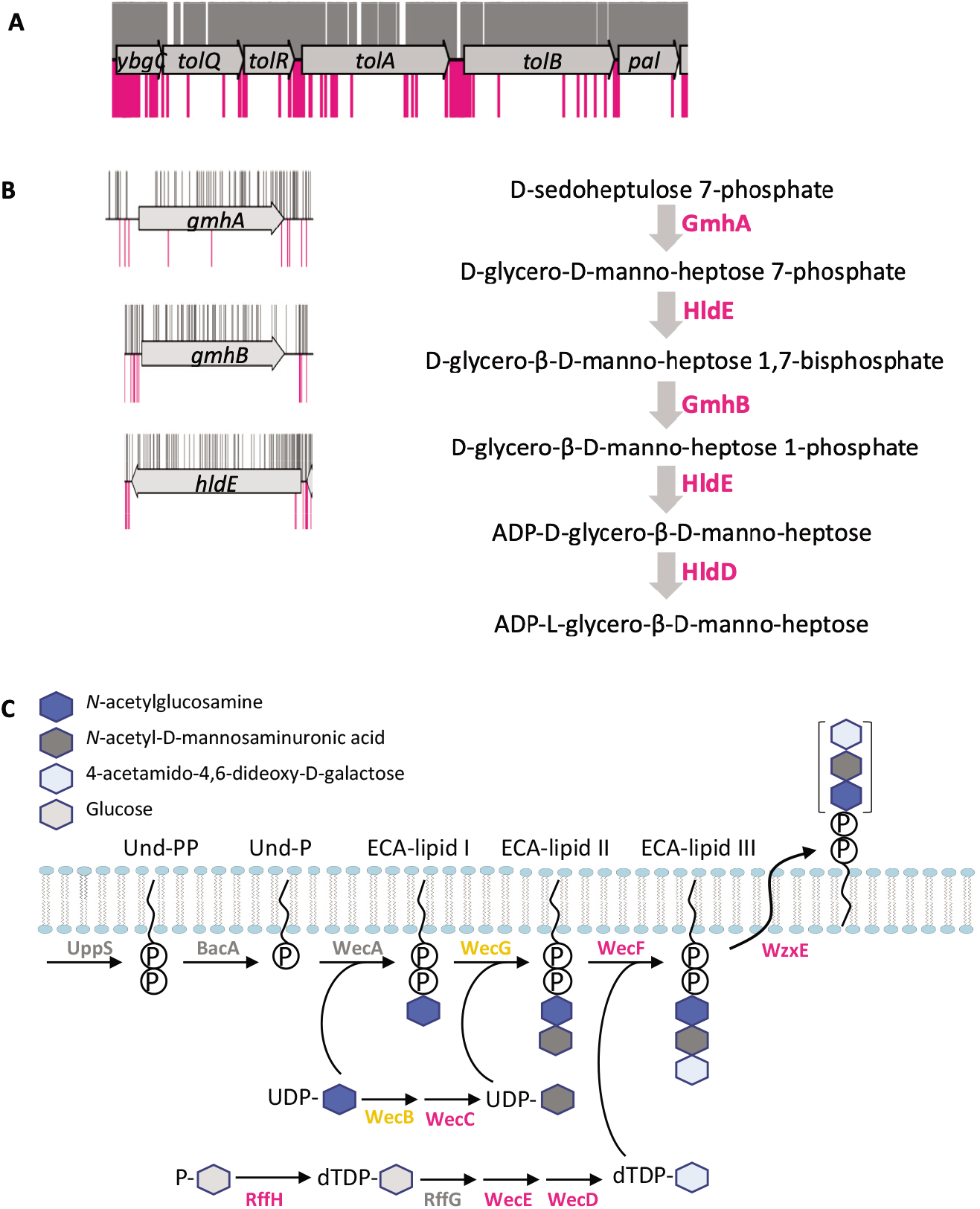
Biosynthetic pathways required in a Δ*yhcB* background. Transposon insertion data of the yhcB library shown in pink, below, control library shown in grey, above, the gene track with all insertion data capped at a frequency of 1. (A) Transposon insertion profile of the *tol-pal* operon. (B) The heptose biosynthetic pathway, required for LPS core biosynthesis. (C) Schematic of the ECA biosynthesis pathway adapted from Jorgenson *et al*. (2016). Abbreviations: Enterobacterial Common Antigen (ECA); Undecaprenyl phosphate (Und-P); Phosphate (P); Adenosine diphosphate (ADP); deoxythymidine diphosphate (dTDP); Uridine diphosphate (UDP).

**Supplementary Figure 8.**
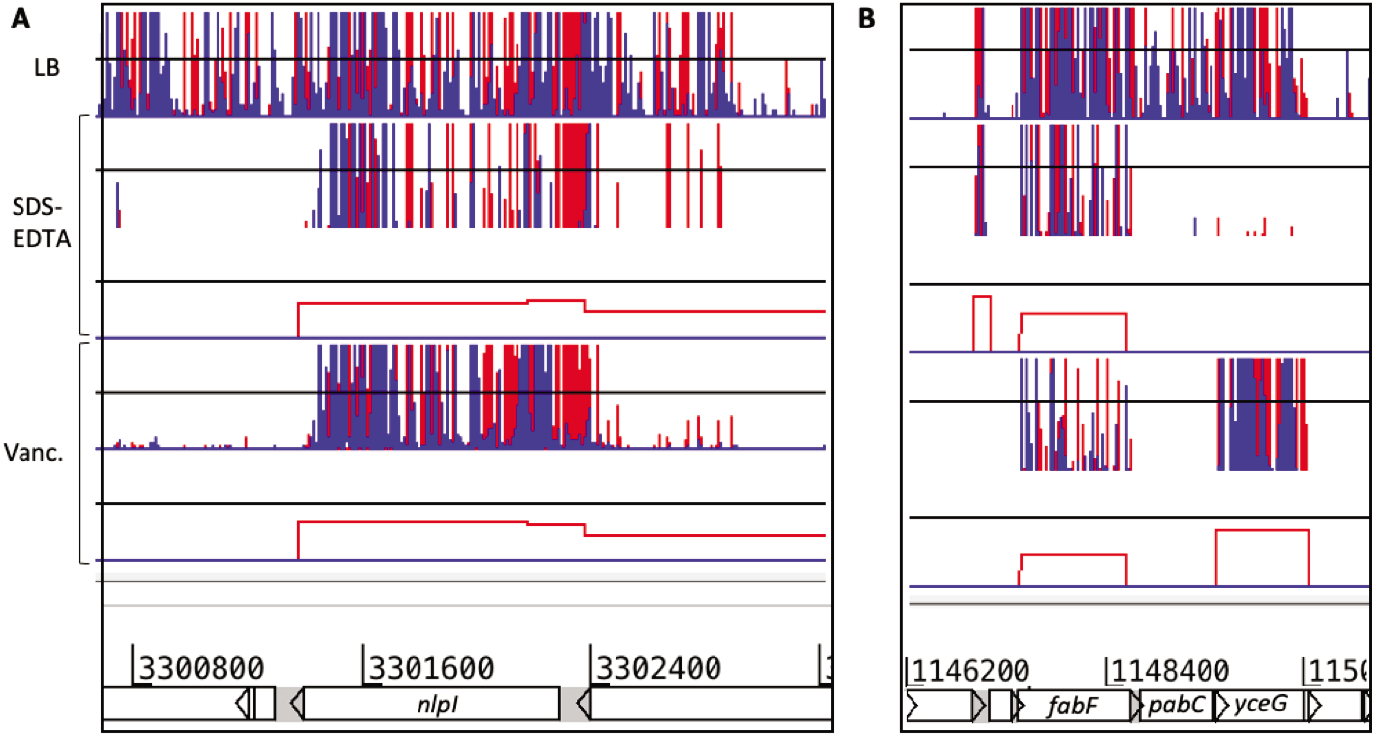
Transposon insertion sites within *nlpl* and *fabF* that restore resistance of a *yhcB* mutant to vancomycin or SDS and EDTA. Transposon insertion data for suppressor screens. Red and blue vertical lines indicate the transposon insertion position and correspond with the transposon orientation at the point of insertion. The height of the bar corresponds with mapped sequencing read frequency. The top track represents data for the library plated on LB. The tracks underneath represent the Δ*yhcB* transposon library plated on LB supplemented with SDS and EDTA, or vancomycin, at lethal doses to the BW25113Δ*yhcB* parent strain. The red boxes underneath each suppressor dataset represent significant differential abundance of insertions in the condition sample compared to the control, identified by AlbaTraDIS.

**Supplementary Figure 9.**
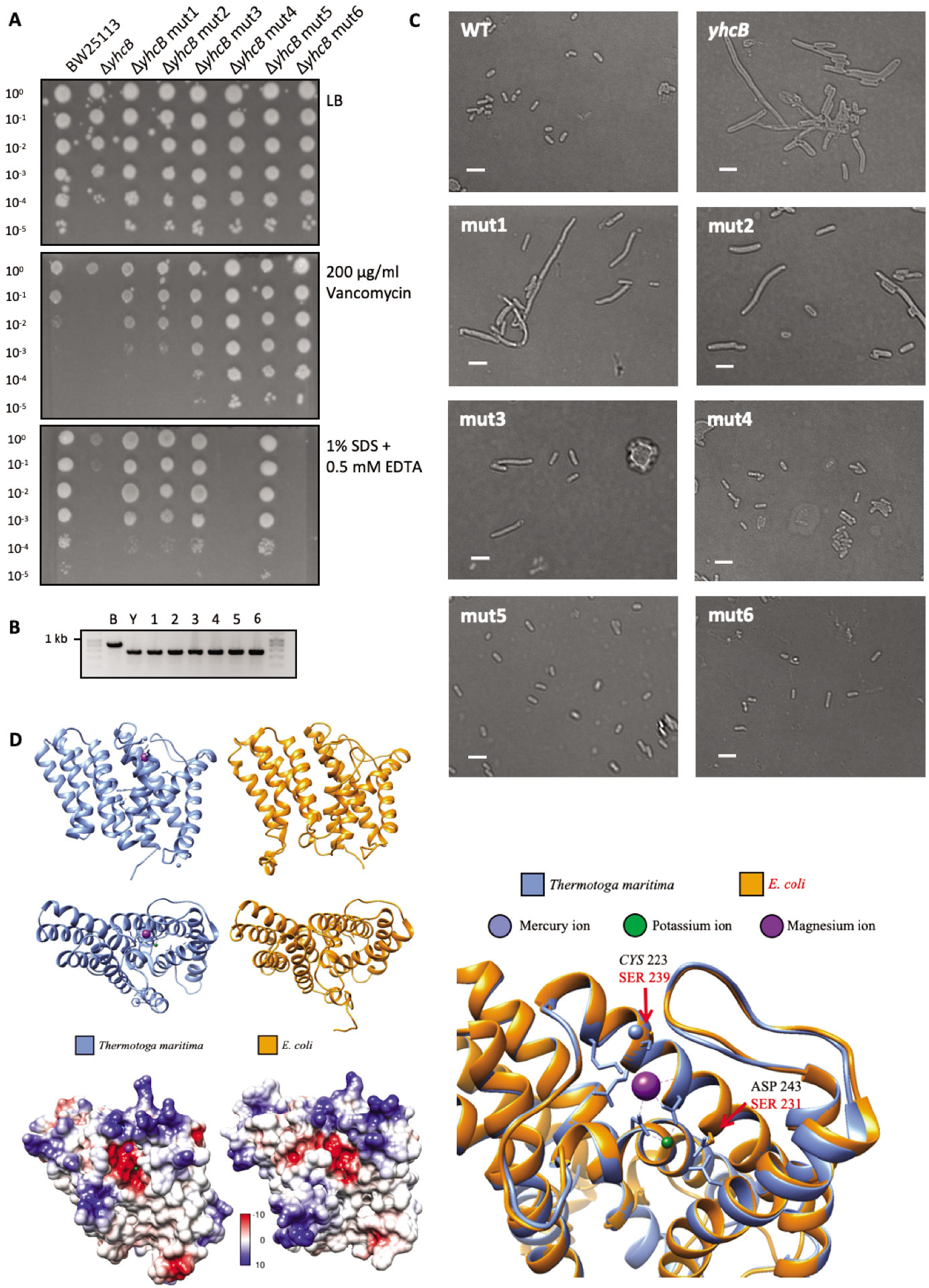
Natural suppressor mutations that are restorative in a *yhcB* mutant. (A) Validation of natural suppressor mutants. 10-fold serial dilutions of E. coli K-12 BW25113, BW25113Δ*yhcB* and six BW25113Δ*yhcB* revertant suppressor mutants, grown on LB, LB supplemented with 200 μg/ml vancomycin, or 1% SDS + 0.5 mM EDTA. (B) PCR amplification of the *yhcB* locus to confirm *yhcB*-deletion in these strains. (C) DIC images of BW25113, BW25113Δ*yhcB* and six BW25113Δ*yhcB* revertant suppressor mutants grown overnight in LB. Scale bar of 5 μm. (D) Solved structure of *Thermatoga maritima* CdsA (blue) and the predicted structure of *E. coli* CdsA (orange), with the surface electrostatic potential of each shown below. The deep red and deep blue colours indicate electronegative and electropositive regions at −10 and 10kT e^−1^, respectively. An overlay of the cation-binding pocket of CdsA from *T. maritima* and *E. coli* is shown in the bottom right panel. The equivalent mutated residues are annotated: shown in black, above for *T. maritima* and red, below for *E. coli*. Note the S223C residue mutated in *T. maritima* to achieve a resolved structure.

**Supplementary Figure 10.**
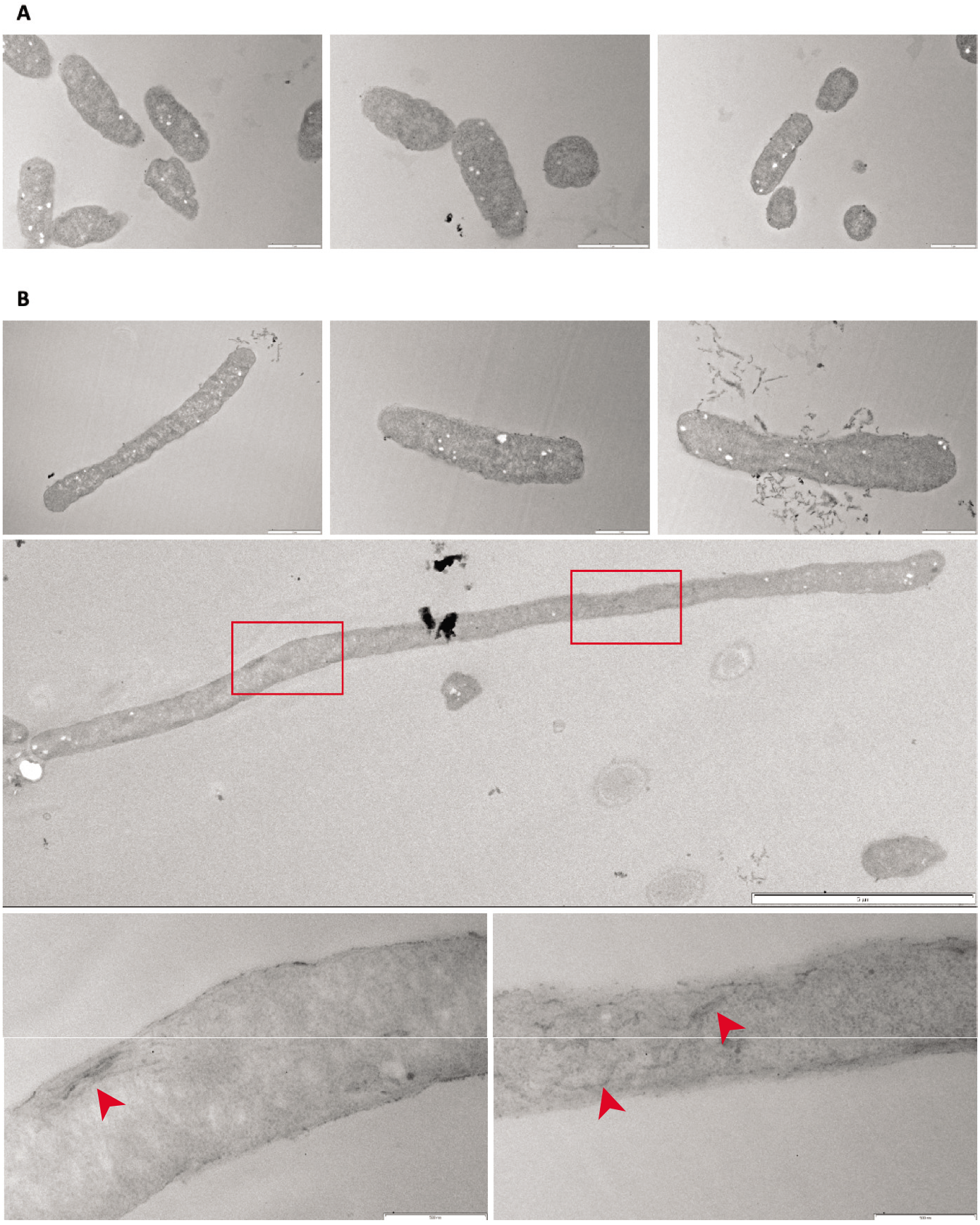
TEM images of BW25113 and the Δ*yhcB* mutant. Transmission electron micrographs of (A) BW25113 and (B) the Δ*yhcB* mutant cells (standard fixation and processing). Scale bar = 1 μm in the top six images. A large Δ*yhcB* mutant cell is shown with a scale bar of 5 μm and two regions of excess or ruffled membrane structures are highlighted by red boxes (annotated in Fig. 10A); enlarged images of these sections are shown underneath with a scale bar of 500 nm each.

**Supplementary Figure 11.**
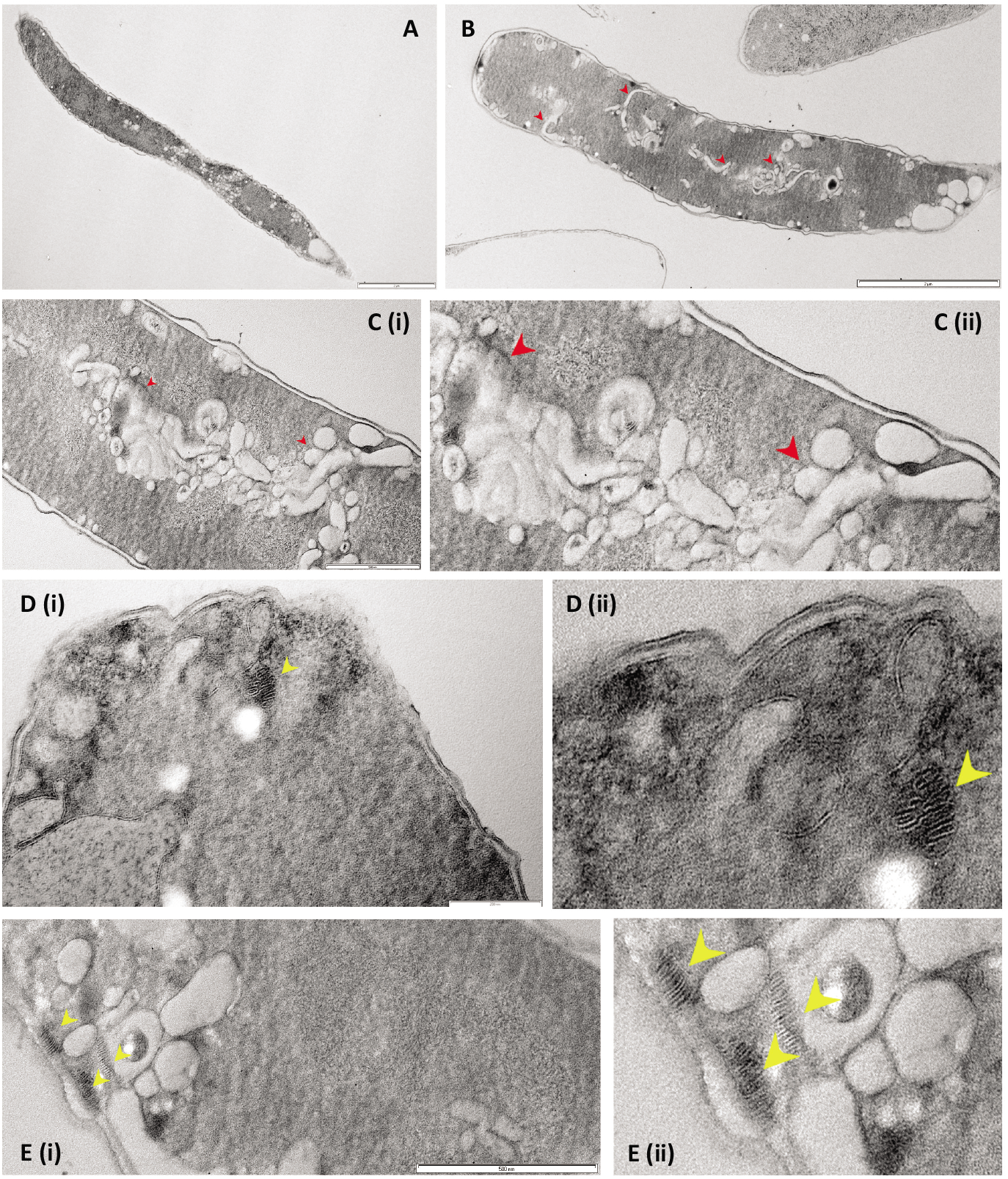
Images of a Δ*yhcB* mutant cells. Electron micrographs of fast frozen/freeze-substituted *yhcB* mutant cells. Internal membranes indicated by red arrowheads; stacked membrane arrays indicated by yellow arrowheads. Bars are: (A) 2 μm; (B) 2 μm; (C) 500 nm; (D) 200 nm; (E) 500 nm. Note that panels C-E (ii) are higher magnification views from C-E (i), respectively.

**Supplementary Figure 12.**
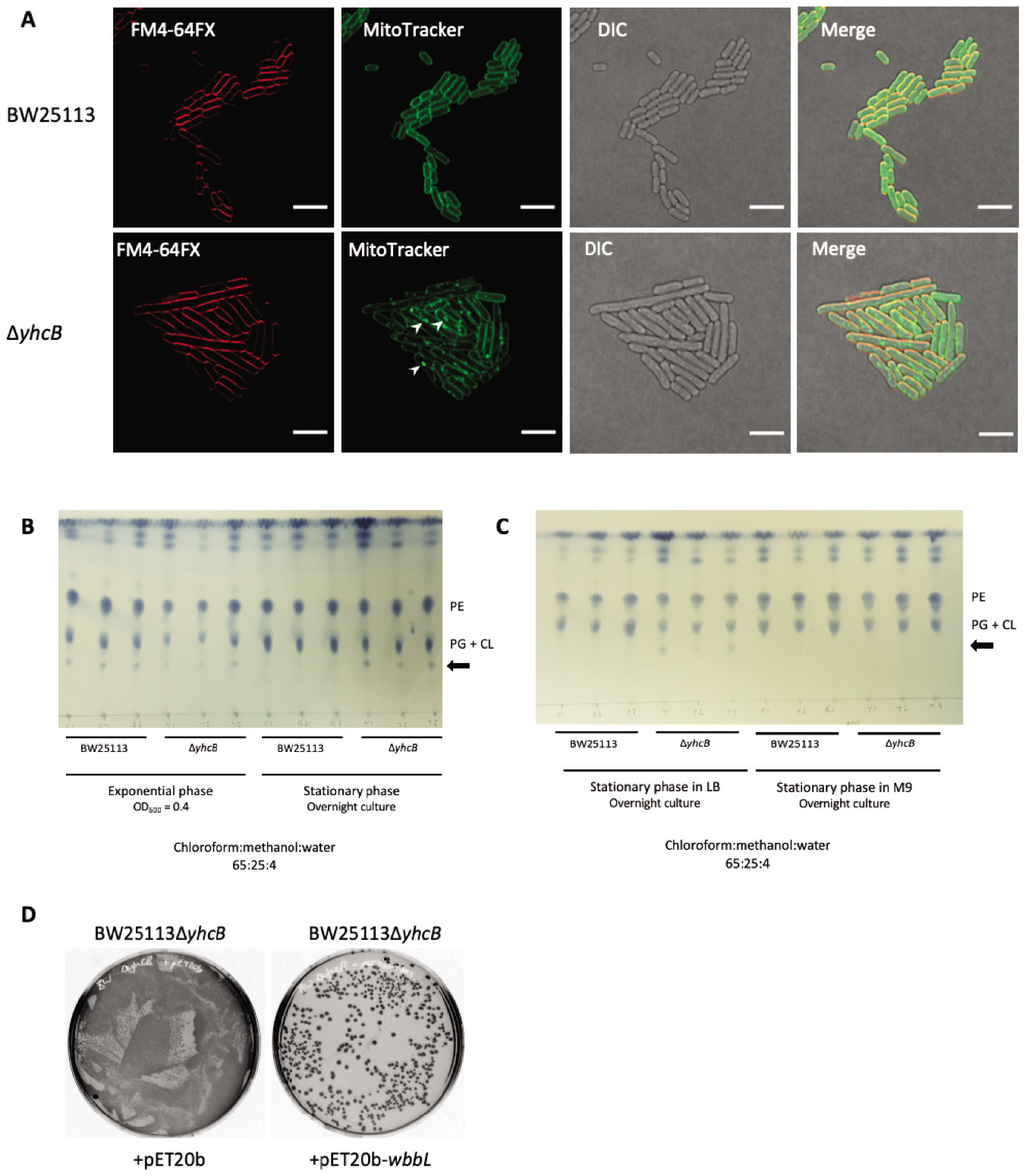
The *yhcB* mutant contains extra membrane, has an altered cell envelope and restoration of O-antigen is toxic. (A) *E. coli* K-12 BW25113 and BW25112Δ*yhcB* grown in LB, labelled with FM4-64FX and MitoTracker Green FM. Images were taken at 3 h and are representative of n = 3 experiments. Lipid accumulation is indicated by the white arrows. Scale bar of 5 μm. (B-C) Thin layer chromatography (TLC) separation of phospholipid species in chloroform:methanol:water (65:25:4) solvent system, and stained with PMA. (B) Samples grown in LB and collected at two stages of growth. (C) Samples grown overnight in LB or M9-glucose. (D) Transformation efficiency of pET20b constructs +/- *wbbL* for restoring O-antigen in a Δ*yhcB* mutant. Abbreviations: Phosphatidylethanolamine (PE); Phosphatidylglycerol (PG); Cardiolipin (CL).

**Supplementary Figure 13.**
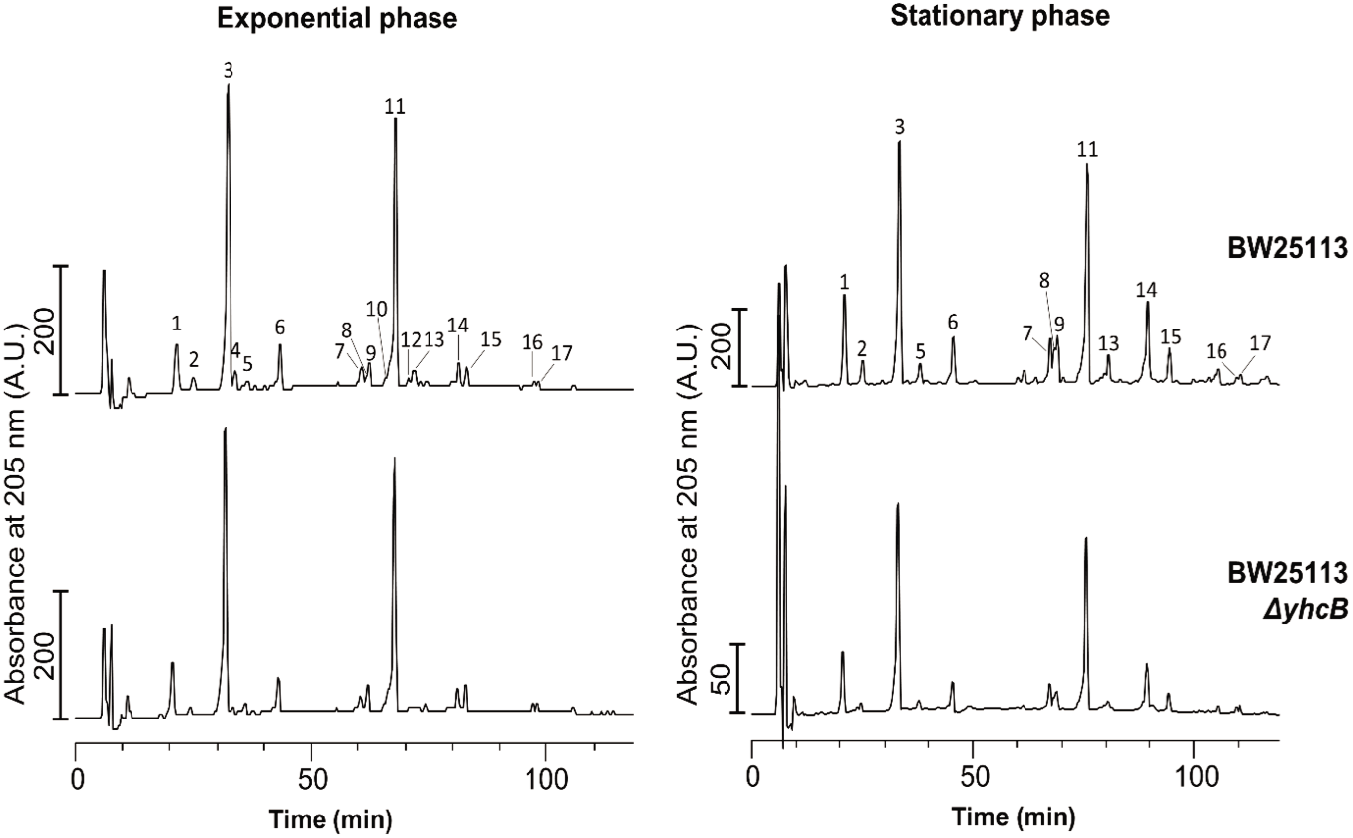
Analysis of the peptidoglycan composition. Representative HPLC chromatograms showing the muropeptide composition of BW25113 and BW25113Δ*yhcB* at exponential and stationary phase. Purified peptdoglycan was digested with cellosyl and the resulting muropeptides were reduced with sodium borohydride and separated by HPLC. The major muropeptides (No. 1-17) are quantified in Table S5.

**Supplementary Figure 14.**
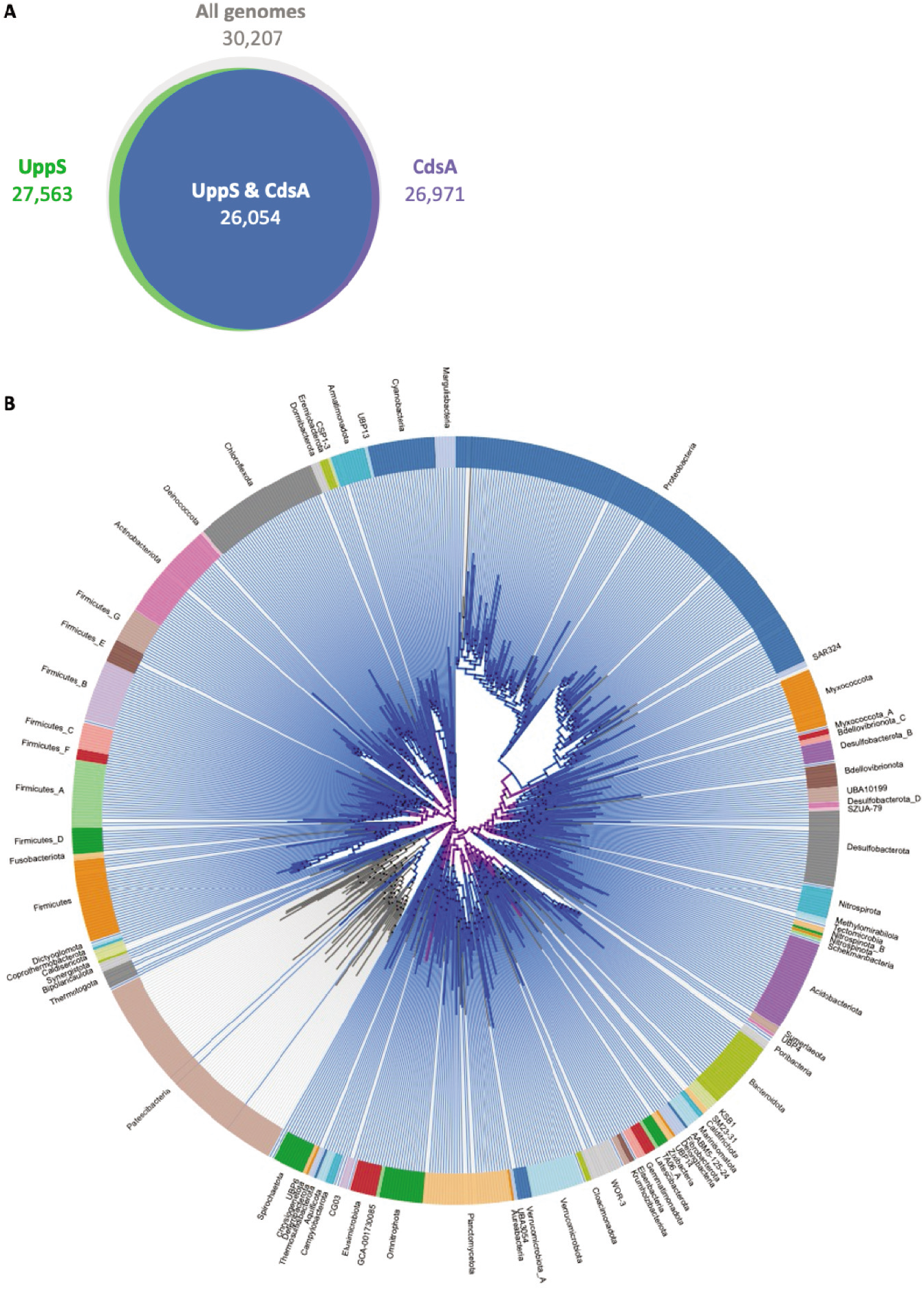
UppS and CdsA conservation. (A) The number, and overlap, of bacterial genomes containing a UppS (K00806) or CdsA (K00981) homolog obtained from AnnoTree (v1.2.0) using the KEGG identifiers and the following search criteria: % identity: 30; E value: 0.00001; % subject alignment: 70; % query alignment: 70. (B) Tree representation of the phylogeny of bacterial genomes with species containing both UppS and CdsA highlighted in blue.

**Supplementary Figure 15.**
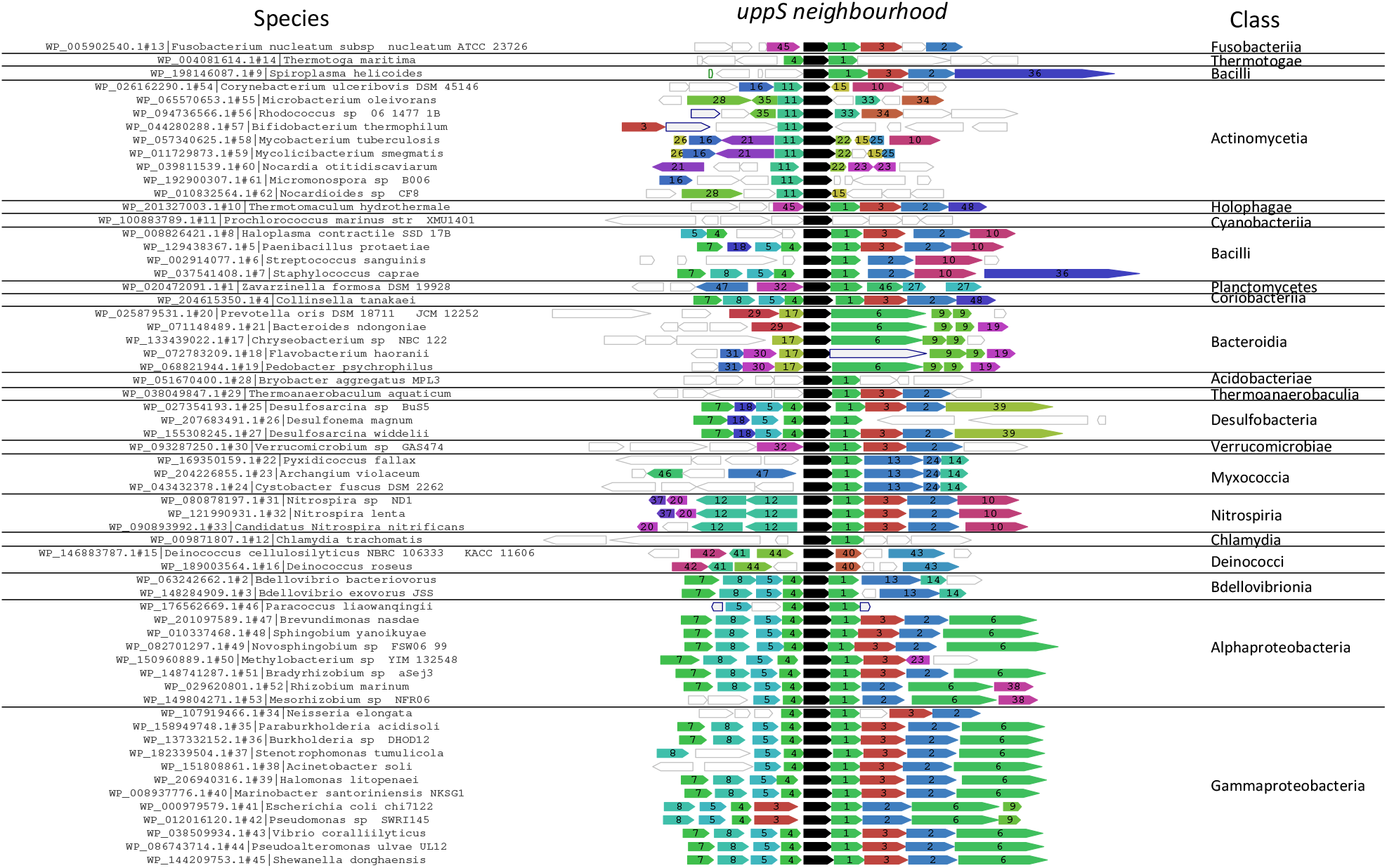
*uppS* and *cdsA* neighbourhood conservation across the bacterial kingdom. The conservation of the *uppS* gene neighbourhood represented by 62 diverse bacterial species. *uppS* is depicted in black, and *cdsA* in green (1). The remaining gene identifiers can be found in Supplementary Table 9. Data calculated using FlaGs (Saha et al. 2020).

## Supplementary Tables

Supplementary Table 1. Polymyxin B TIS screen data

Supplementary Table 2. Transposon library construction metrics

Supplementary Table 3. Insertion index scores of the *yhcB* and WT transposon libraries

Supplementary Table 4. Gene fitness in a *yhcB* mutant strain

Supplementary Table 5. Peptidoglycan composition

Supplementary Table 6. PANTHER analysis

Supplementary Table 7. AlbaTraDIS identification of suppressor mutations

Supplementary Table 8. Mutations identified in spontaneous revertant suppressor mutants

Supplementary Table 9. FlaGs gene identifiers

